# Genome-wide signatures of plastid-nuclear coevolution point to repeated perturbations of plastid proteostasis systems across angiosperms

**DOI:** 10.1101/2020.08.28.272872

**Authors:** Evan S. Forsythe, Alissa M. Williams, Daniel B. Sloan

## Abstract

Nuclear and plastid (chloroplast) genomes experience different mutation rates, levels of selection, and transmission modes, yet key cellular functions depend on coordinated interactions between proteins encoded in both genomes. Functionally related proteins often show correlated changes in rates of sequence evolution across a phylogeny (evolutionary rate covariation or ERC), offering a means to detect previously unidentified suites of coevolving and cofunctional genes. We performed phylogenomic analyses across angiosperm diversity, scanning the nuclear genome for genes that exhibit ERC with plastid genes. As expected, the strongest hits are highly enriched for plastid-targeted proteins, providing evidence that cytonuclear interactions affect rates of molecular evolution at genome-wide scales. Many identified nuclear genes function in post-transcriptional regulation and the maintenance of protein homeostasis (proteostasis), including protein translation (in both the plastid and cytosol), import, quality control and turnover. We also identified nuclear genes that exhibit strong signatures of coevolution with the plastid genome but lack organellar-targeting annotations, making them candidates for having previously undescribed roles in plastids. In sum, our genome-wide analyses reveal that plastid-nuclear coevolution extends beyond the intimate molecular interactions within chloroplast enzyme complexes and may be driven by frequent rewiring of the machinery responsible for maintenance of plastid proteostasis in angiosperms.

## Introduction

Only a small fraction of the proteins required for plastid function are encoded by the plastid genome (plastome) itself (Timmis et al., 2004; van Wijk & Baginsky, 2011). The remaining plastid-localized proteins are encoded in the nuclear genome, translated in the cytosol, and imported into plastids (hereafter referred to as nuclear-encoded plastid-targeted [N-pt] proteins), where they often interact with the plastome and its gene products (Gould et al., 2008). These plastid-nuclear interactions are critical for overall fitness, as evidenced by the frequent role of plastid-nuclear incompatibilities in reproductive isolation (Schmitz-Linneweber et al., 2005; Greiner et al., 2011; Bogdanova et al., 2015; Barnard-Kubow et al., 2016; Zupoka et al., 2020).

One signature of proteins that are functionally related and/or coevolving is that they tend to exhibit correlated changes in rates of sequence evolution across a phylogeny, which is known as evolutionary rate covariation (ERC) and can be quantified by comparing genetic distances or branch lengths of gene trees from two potentially interacting genes (Goh et al., 2000; Ramani & Marcotte, 2003; Sato et al., 2005; Nathaniel L. Clark & Aquadro, 2010; Nathan L Clark et al., 2012; De Juan et al., 2013). The known physical interactions within “chimeric” plastid-nuclear complexes (i.e., those containing both plastome-encoded and N-pt proteins) have provided a valuable system to test and illustrate the principle that coevolution and functional interactions can result in ERC (Sloan, Triant, Wu, et al., 2014; Zhang et al., 2015, 2016; Rockenbach et al., 2016; Weng et al., 2016; Williams et al., 2019).

In addition to probing known interactions, ERC has served as a powerful tool to scan entire genomes/proteomes to detect previously unrecognized functional relationships (Findlay et al., 2014; Raza et al., 2019), which do not always entail direct physical interactions (Nathan L Clark et al., 2012). For example, application of a genome-wide ERC scan in diverse insects with heterogeneous rates of mitochondrial genome evolution recovered novel mitonuclear interactions (Yan et al., 2019). However, despite strong evidence of correlated rates among known members of plastid-nuclear complexes, ERC analysis has not been applied on a genome-wide scale across diverse plant lineages, meaning we may have only scratched the surface with respect to the full breadth of plastid-nuclear interactions. A key barrier is that the frequent occurrence of gene and whole-genome duplication in plants (Panchy et al., 2016; Wendel et al., 2018) makes it inherently difficult to perform phylogenomic scans for ERC. Typical implementations of ERC analysis require one-to-one orthology in gene trees (Nathan L Clark et al., 2012; Findlay et al., 2014; Wolfe & Clark, 2015; Yan et al., 2019), but gene duplication yields large gene families composed of sequences that share both orthology and paralogy (Bansal & Eulenstein, 2008; Stolzer et al., 2012). Outside of the context of ERC, numerous studies have overcome challenges associated with phylogenomics in plants by carefully filtering gene families and/or extracting subtrees that represent mostly orthologs (Sanderson & McMahon, 2007; Duarte et al., 2010; De Smet et al., 2013; Sangiovanni et al., 2013; Forsythe et al., 2020). Nevertheless, these approaches cannot completely eliminate the pervasive effects of gene duplication and differential loss, so performing ERC analyses across diverse plant lineages requires a novel approach that can accommodate this recurring history.

ERC analyses have the potential to be especially powerful for probing plastid-nuclear interactions because the rate of plastome evolution can differ greatly across angiosperm species, with several lineages exhibiting extreme accelerations. Not surprisingly, angiosperms that lose photosynthetic function and transition to parasitic/heterotrophic lifestyles exhibit massive plastome decay and rapid protein sequence evolution (Wicke et al., 2016), in extreme cases resulting in outright loss of the entire plastome (Molina et al., 2014). However, even among angiosperms that remain fully photosynthetic, there have been repeated accelerations in rates of plastid gene evolution (Jansen et al., 2007; Guisinger et al., 2008; Knox, 2014; Sloan, Triant, Forrester, et al., 2014; Dugas et al., 2015; Nevill et al., 2019; Shrestha et al., 2019). These accelerations in angiosperms that retain a photosynthetic lifestyle can be highly gene-specific (Magee et al., 2010) and are often most pronounced in non-photosynthetic genes, such as those that encode ribosomal proteins, RNA polymerase subunits, the plastid caseinolytic protease (Clp) subunit ClpP1, the acetyl-CoA carboxylase (ACCase) subunit AccD, and the essential chloroplast factors Ycf1 and Ycf2 (Guisinger et al., 2008; Sloan, Triant, Forrester, et al., 2014; Seongjun Park et al., 2017; Shrestha et al., 2019). Accelerated protein sequence evolution has frequently been accompanied by other forms of plastome instability, including structural rearrangements and gene duplication (Guisinger et al., 2011; Knox, 2014; Sloan, Triant, Forrester, et al., 2014; Shrestha et al., 2019) as well as accelerated mitochondrial genome evolution in some cases (Cho et al., 2004; Parkinson et al., 2005; Jansen et al., 2007; Mower et al., 2007; Sloan et al., 2009; Seongjun Park et al., 2017). Several explanations have been proposed for the cause of these cases of rapid plastome evolution, but they remain largely a mystery (Guisinger et al., 2008; Seongjun Park et al., 2017; Williams et al., 2019). Discovering the full suite of nuclear genes that repeatedly co-accelerate with plastid genes may advance our understanding of this angiosperm evolutionary puzzle.

Here, we develop an approach to apply genome-wide ERC analyses across diverse angiosperms to identify hundreds of nuclear genes that exhibit signatures of ERC with the plastome. This set of genes is highly enriched for known N-pt genes with functions in several pathways that appear to be centered around maintenance of plastid protein homeostasis (proteostasis). We also observe strong signatures of plastid-nuclear ERC for more than 30 non-plastid-targeted proteins, representing candidates for novel plastid-nuclear interactions. Together, our findings impact our understanding of the genome-wide landscape of plastid-nuclear interactions.

## Results

### Genome-wide ERC analyses detect correlated evolution between the plastome and N-pt genes

To perform a genome-wide scan for plastid-nuclear ERC, we partitioned the plastid-encoded proteins from 20 angiosperms known to exhibit a wide range of plastome evolutionary rates (Fig. 1) into seven functional categories: AccD, ClpP1, MatK, photosynthesis, ribosomal proteins, RNA polymerase, and Ycf1/Ycf2 (Table S1). We applied a custom phylogenomic analysis pipeline to a matching set of nuclear genomes and transcriptomes (Fig. 2). Our pipeline included steps designed to extract gene families sharing orthology in the presence of gene duplication and loss. It yielded a filtered set of 7929 gene trees with an average of 25.1 sequences per tree and 16.4 species per tree (Fig. S1). Our genome-wide scan for plastid-nuclear ERC was executed by testing all possible 55,503 pairwise correlations between trees (7 plastome trees x 7929 nuclear trees) based on normalized branch lengths to account for lineage-specific features that may affect rates across entire genomes (e.g., generation time) (Nathaniel L. Clark & Aquadro, 2010). To directly compare trees that can differ in topology, gene duplication, and species representation, we measured branch lengths for each species on each tree using a ‘root-to-tip’ approach (Yan et al., 2019), in which we averaged the cumulative branch length of the path leading from the common ancestor of all monocots and eudicots to each tip (gene copy) for each species (see Methods).

**Figure 1.**
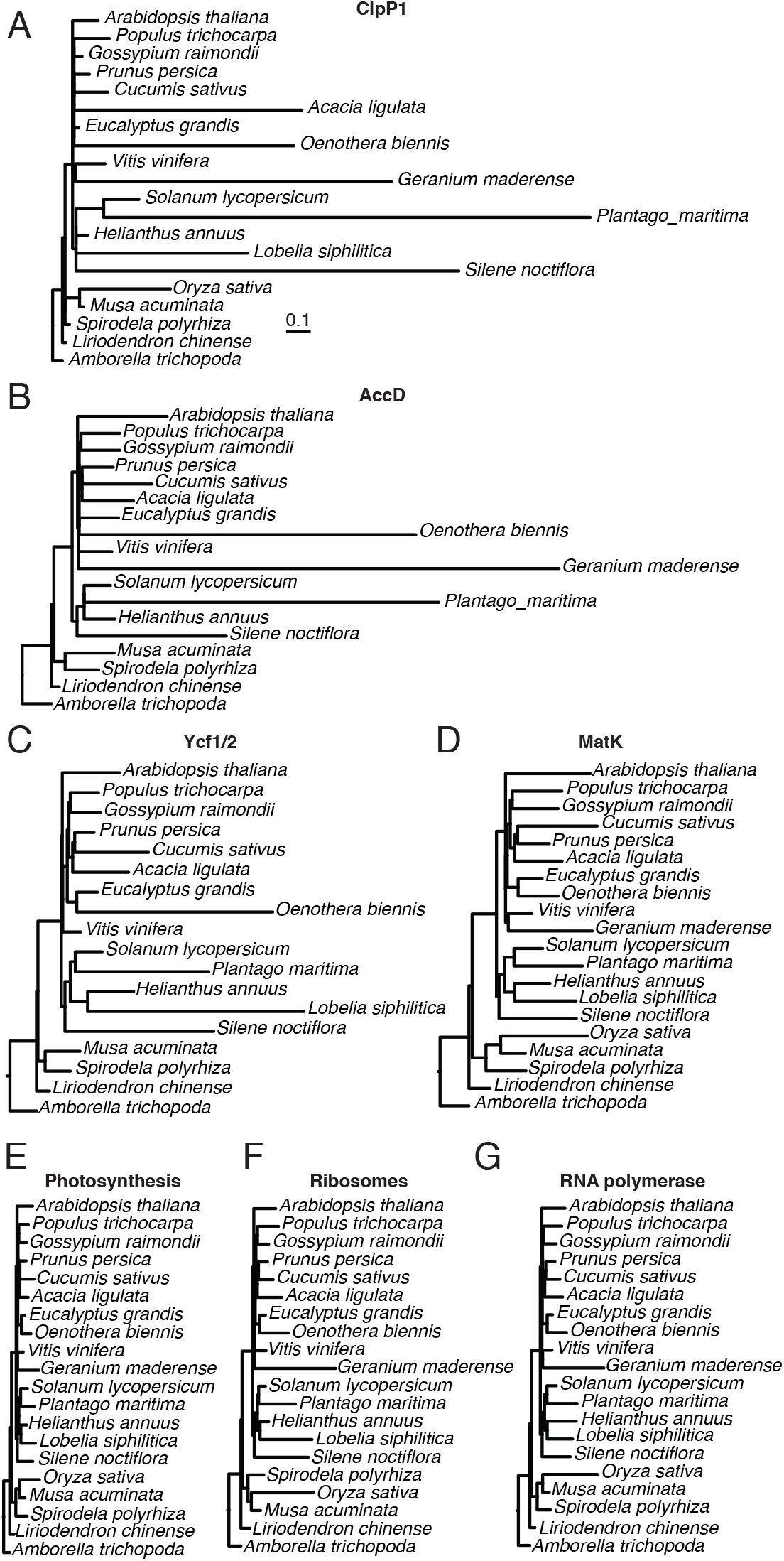
Trees based on plastome partitions. Branch length optimized trees inferred from amino acid sequence alignments for plastid genes partitioned into seven functional categories (described in Table S1). Branch lengths are shown on the same scale for all trees to highlight differences in rates of amino acid evolution among partitions. Each plastome partition tree was used for ERC analysis against all nuclear gene trees.

**Figure 2.**
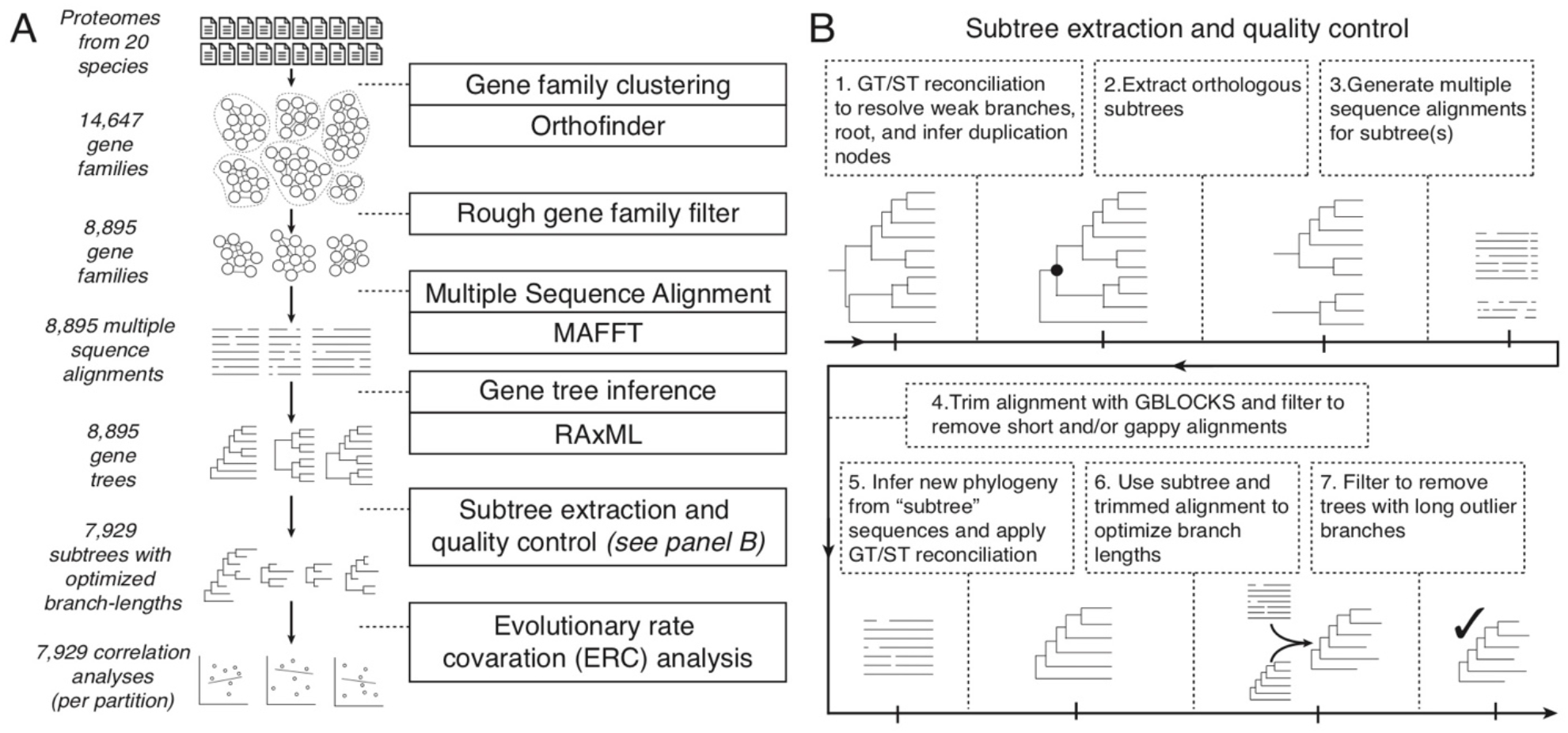
Phylogenomic pipeline used to identify and analyze nuclear gene families. (A) Flowchart depicting the steps leading up to ERC analyses. (B) Steps of the extraction and quality-control procedure.

To illustrate the ERC principle, we highlight a case study from the plastid Clp complex, which is composed of the plastid-encoded ClpP1 subunit and multiple N-pt subunits (Nishimura & van Wijk, 2015). This complex represents an effective positive control in the context of a genome-wide scan because it was previously shown to exhibit strong ERC signals among subunits (Rockenbach et al., 2016; Williams et al., 2019). The Clp complex core is composed of two heptameric rings, the ‘R-ring’ and ‘P-ring’. ClpP1 is part of the R-ring and interacts more closely with the other subunits in this ring (ClpR subunits) than with the subunits of the P-ring (ClpP subunits) (Nishimura & van Wijk, 2015). These core rings are also accompanied by a variety of accessory proteins (ClpC, ClpD, ClpF, ClpS, and ClpT subunits), allowing us to compare ERC results for N-pt genes with varying degrees of physical interaction. A mirrored tree diagram of ClpP1 and ClpR1 illustrates that branch lengths from corresponding species on the two trees exhibit strong ERC (R^2^ = 0.94; Fig. 3A-B). Extending this analysis to all nuclear genes, a genome-wide distribution of ERC results for ClpP1 reveals that all ClpR and ClpP subunits are present among the strongest ERC hits (top 2% of all genes analyzed), and all but one of these maintains genome-wide significance after correcting for multiple tests (Fig. 3C). Further, we find a general pattern of clustering of ERC values between ClpP1 and other Clp subunits that corresponds to the intimacy of their known interactions; ClpR subunits display the strongest ERC, followed by ClpP subunits, with the accessory Clp subunits showing the weakest signal. Thus, ERC appears to be sufficiently sensitive to detect functional plastid-nuclear interactions even with the background of a genome-wide scan.

**Figure 3.**
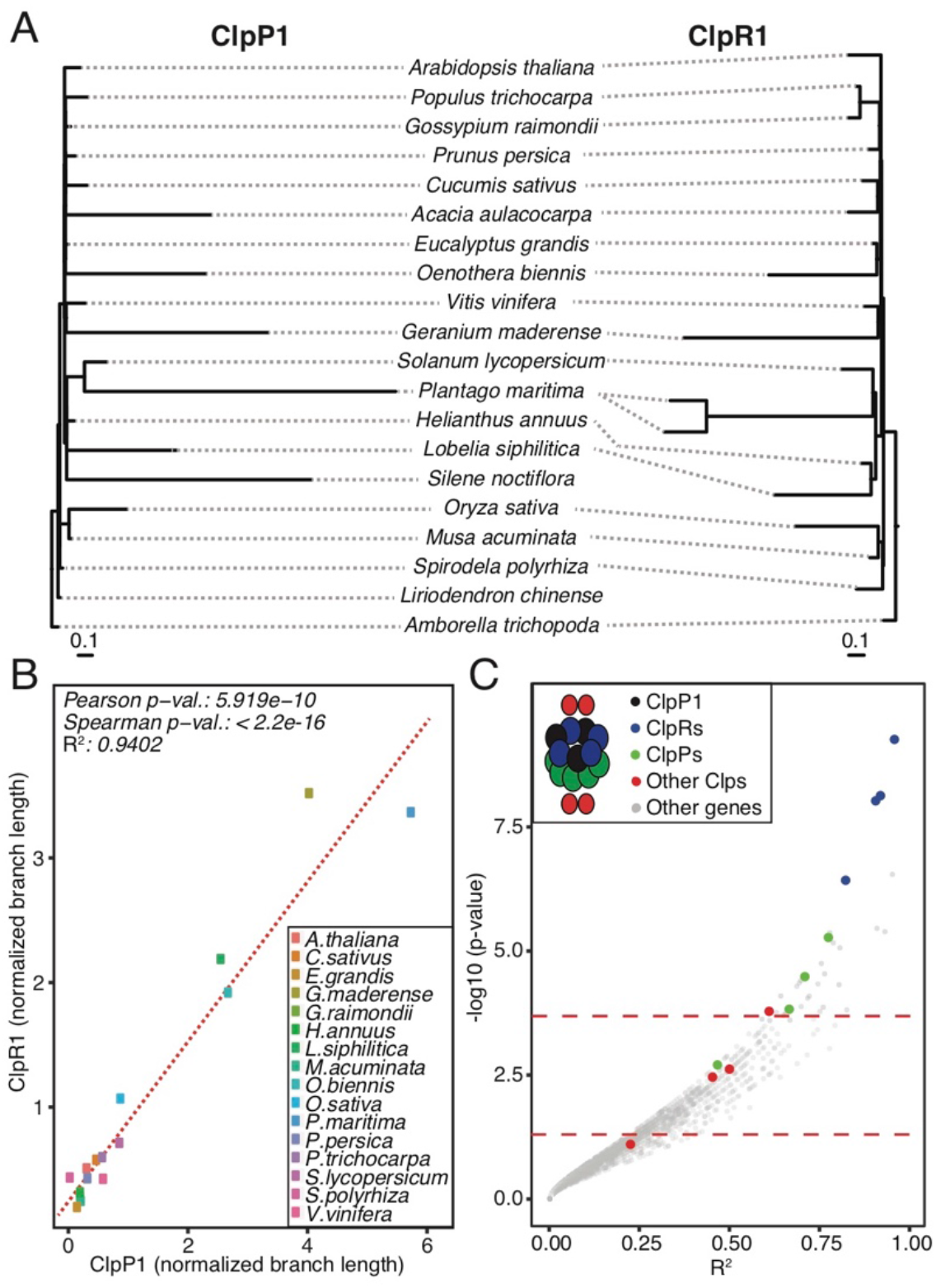
Case-study of ERC between plastid-encoded ClpP1 and nuclear gene trees. (A) ClpP1 and ClpR1 gene trees shown mirrored to highlight correlation of branch lengths. (B) Linear regression quantifying correlation of evolutionary rates between ClpP1 and ClpR1. Points represent normalized branch lengths estimated from ClpP1 (x-axis) and ClpR1 (y-axis) gene trees. Red dotted line indicates best fit trend line. (C) Results from ERC analyses of ClpP1 versus all nuclear genes. Each point represents p-value and R^2^ values from a pairwise ERC analysis (Pearson correlation). ERC comparisons with negative slopes are not shown. Known Clp complex nuclear genes are colored by their placement in the Clp structure (depicted in the legend). Red dotted line indicates a raw p-value of 0.05 (bottom) and a genome-wide significance at an FDR-corrected p-value of 0.05 (top).

We performed ERC analyses in parallel for each of the seven plastome partition trees against normalized branch lengths from the nuclear trees (Table S2). We found that N-pt genes are highly significantly overrepresented in ERC hits for all plastome partitions, displaying roughly two-fold enrichment (Fig. 4). We identified the subset of these genes that are known to directly physically interact with plastid-encoded proteins (Forsythe et al., 2019) and observed an even higher degree of enrichment (approximately 4-fold to 8-fold depending on the plastome partition). We also found correlations between plastome partitions and nuclear genes with mitochondrial function. Overall, mitochondrial-targeted (N-mt) proteins are significantly enriched among ERC hits for all plastome partitions except for RNA polymerase and photosynthesis, although the effect size (approximately 1.5-fold) was smaller than for N-pt genes. N-mt proteins involved in direct physical interactions with mitochondrial-encoded proteins showed an increased degree of enrichment compared to all N-mt proteins (approximately two-fold), which was significant for all partitions. Proteins with dual localization to both plastids and mitochondria displayed wider variance of enrichment with inconsistent significance, both of which may be related to the small sample size of this gene category. Finally, we found that genes annotated as localized to any parts of the cell other than the plastids or mitochondria are significantly depleted among ERC hits for all partitions (Fig. 4). These results indicate that correlated plastid-nuclear evolution is pervasive across the nuclear genomes and this signature is detectable by ERC.

**Figure 4.**
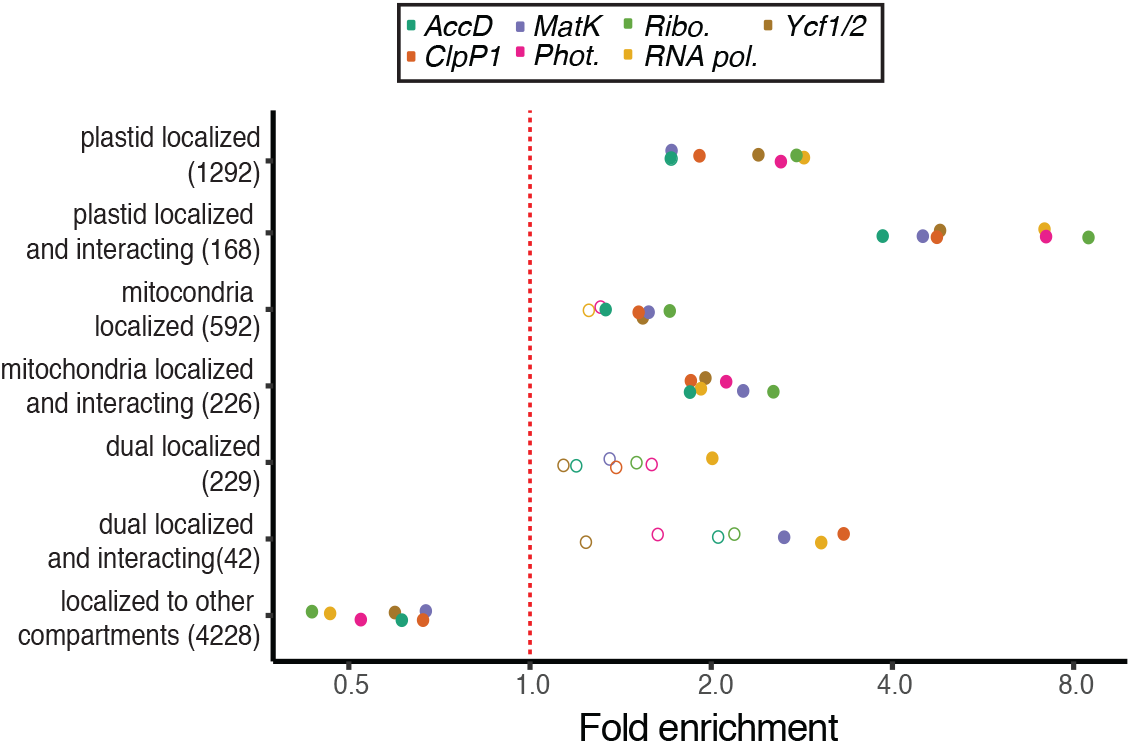
Subcellular localization and cytonuclear interactions of ERC hits. Genes exhibiting signatures of coevolution with plastome partitions were analyzed for their localization and interactions as classified by the *CyMIRA* database (Forsythe et al., 2019). Categories indicating ‘interacting’ refers to nuclear proteins predicted to directly physically interact with organelle-encoded proteins. The number of total genes in each category are indicated in parentheses. Statistical significance of enrichment/depletion (Fisher’s exact test) is indicated by filled points (p < 0.05).

### Functions associated with plastid proteostasis are highly enriched in ERC hits

Gene Ontology (GO) analyses of the ERC hits showed that several categories associated with plastid and mitochondrial function were significantly enriched, while GO terms associated with other cellular compartments (e.g., ‘Nuclear’ and ‘Endomembrane’) were significantly depleted (Fig. 5). Combined with the targeting data presented above (Fig. 4), these results reinforce the power of ERC in detecting cytonuclear interactions. Further, many of the enriched GO terms are more specifically connected to regulation of plastid proteostasis (Fig. 5). For example, terms related to proteolytic activity (e.g. ‘protein quality control’, ‘chloroplastic Clp complex’, and ‘peptidase activity’) display some of the highest degree of enrichment (more than 8-fold in some cases). This signature is further supported by detection of multiple subunits related to FtsH metalloproteases (Table 1). Translational machinery is also prominent; we found enrichment for several related GO categories (e.g. ‘translation’, ‘ribosome biogenesis’, ‘chloroplast rRNA processing’), and many individual genes that encode plastid ribosomal proteins or are involved in translation initiation/elongation (Table 1). The GO terms ‘protein transmembrane transport’ and ‘protein localization to chloroplast’ are also enriched, indicating genes involved in chloroplast protein import (Table 1). The above functions constitute key regulators of plastid proteostasis (Kim et al., 2013; Dogra et al., 2019), pointing to a possible driver of plastid-nuclear coevolution.

**Table 1:**
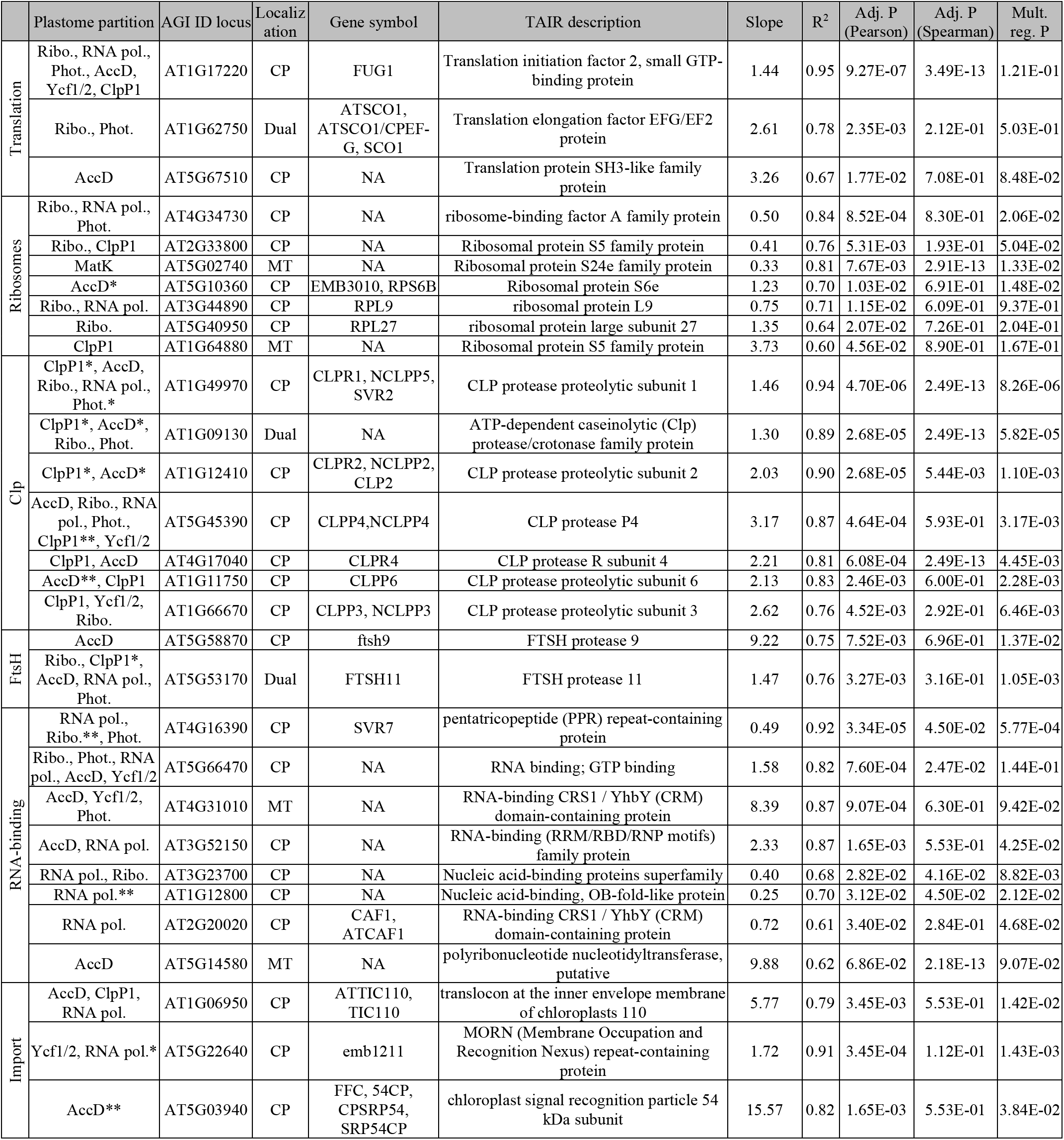
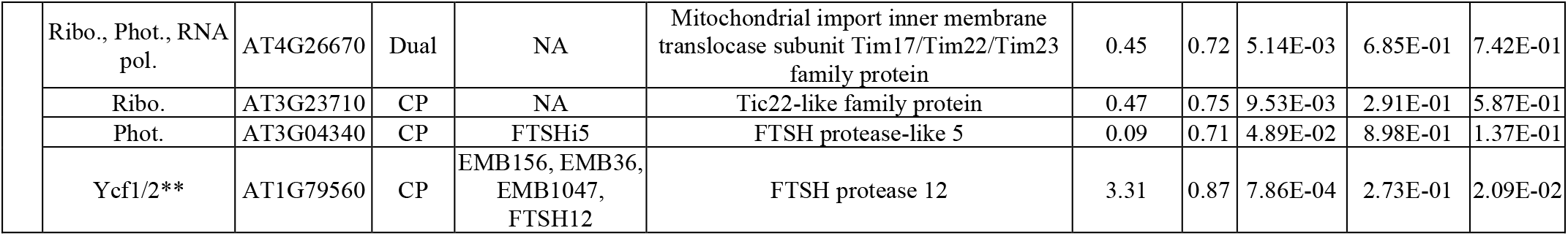
Organelle-localized strong ERC hits. AGI locus identifiers are shown for nuclear genes with significant ERC with plastome partition(s). * indicates significant ERC for the partition in multiple regression. ** indicates the shown partition was the only significant ERC under multiple regression. For genes that are hits in multiple plastome partitions, the slope, R^2^, and P-values for partition with the lowest Pearson P-value are reported. Shown here is a subset of the 99 total organelle-localized strong ERC hits. For full results see Supplementary Data.

**Figure 5.**
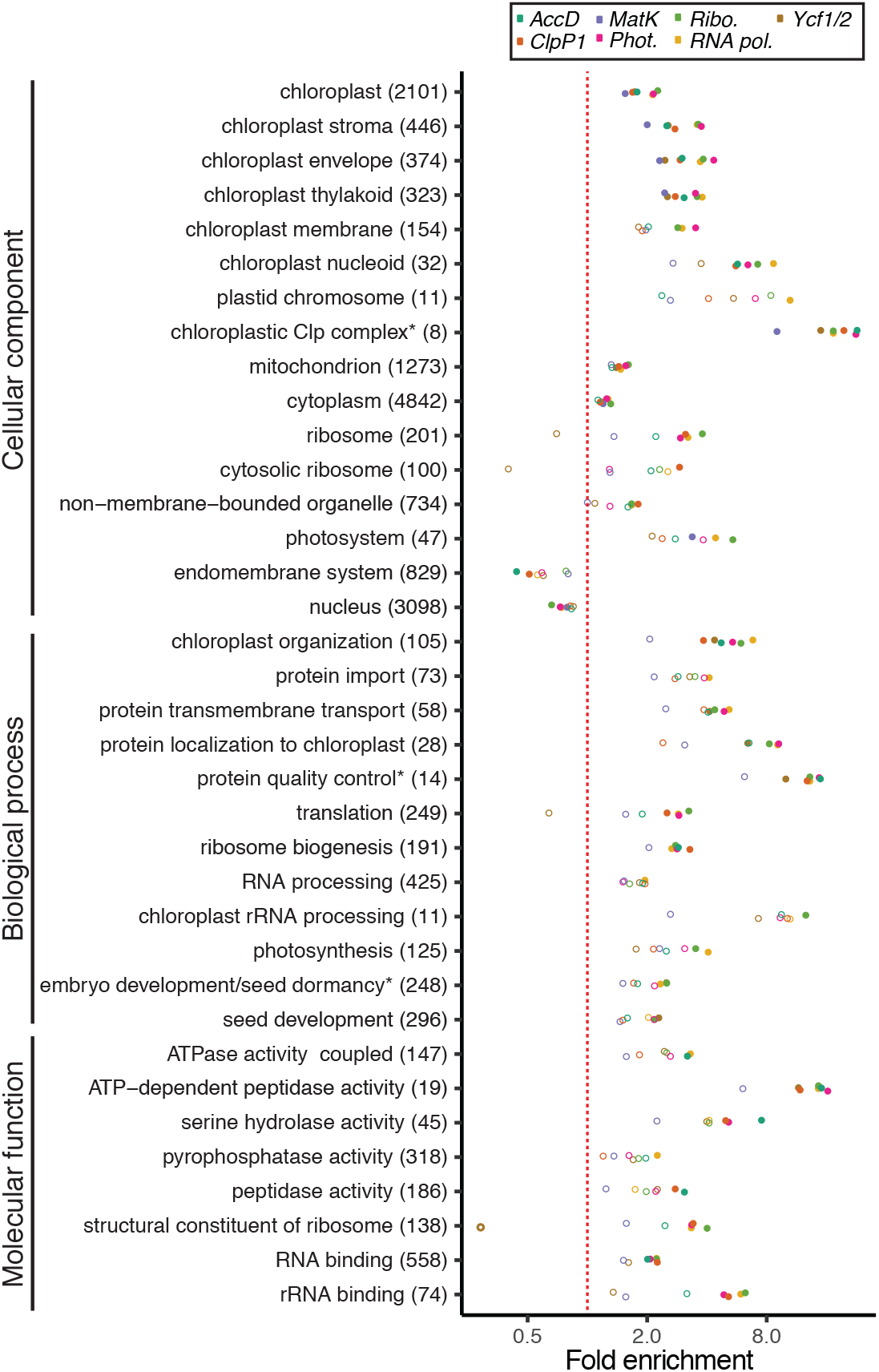
Functional enrichment of ERC hits. Gene Ontology (GO) functional enrichment analyses were performed for ERC hits from each of the plastome partitions. Categories with significant enrichment/depletion in at least one partition are shown. Categories are grouped by type of GO annotation (cellular component, biological process, molecular function). Some redundant or highly overlapping categories were removed (see Supplementary Data for full results). Asterisks indicate shortening of category name to fit figure dimensions. The number of total genes in each category are indicated in parentheses. Statistical significance of enrichment/depletion (Fisher’s exact test) is indicated by filled points (p < 0.05). P-values were corrected for multiple tests using FDR.

Interestingly, the only significantly enriched GO category that is not directly related to plastid or mitochondrial-localized function was ‘cytosolic ribosome’, which also has a clear role in translation. We found that each of the identified cytosolic ribosome gene families contained multiple *Arabidopsis* paralogs, and we confirmed that these were *bona fide* cytosolic ribosomal subunits rather than misannotations of plastid ribosomal subunits in the GO classification scheme (Fig. S3). This result suggests that factors that impact the rate of evolution of plastid genes (and N-pt interaction partners) may also impact cytosolic ribosomes, pointing to potential regulation of plastid proteostasis via maintenance of cytonuclear stoichiometry (see Discussion).

### ERC analyses identify candidates for novel plastid functions

As previously mentioned, the individual hits with the strongest signatures of ERC are dominated by known N-pt or N-mt genes (76%; Table 1). These hits include eleven genes that have been annotated as organelle-localized but designated as ‘proteins of unknown function’. ERC for these genes provides evidence that could help resolve their roles in plastids. In addition, we observed 31 genes (24%) that are not annotated as plastid or mitochondrial-localized (Table 2). These are candidates for novel N-pt genes and may contribute to some of the functions described in the previous section. We discuss some of the most intriguing examples below, including potential novel plastid proteostasis regulators. In sum, our results indicate the specific pathways that exhibit plastid-nuclear ERC and reveal novel N-pt candidates, leading to new hypotheses to advance our understanding of the full scope of plastid-nuclear interactions and their impact on plant evolution.

**Table 2:** Strong ERC hits lacking organelle-localized annotation. AGI locus identifiers are shown for nuclear genes with significant ERC with plastome partition(s). * indicates significant ERC for the partition in multiple regression. ** indicates the shown partition was the only significant ERC under multiple regression. For genes that are hits in multiple plastome partitions, the Slope, R^2^, and P-values for partition with the lowest Pearson P-value are reported. *O. sativa* IDs are shown for families in which *A. thaliana* is not present. One ERC hit lacking an *A. thaliana* and *O. sativa* ID was omitted. For full results see Supplementary Data.

| Plastome partition | AGI locus | Gene symbol | TAIR description | Slope | R<sup>2</sup> | Adj. P (Pearson) | Adj. P (Spearman) | Mult. reg. P |
| --- | --- | --- | --- | --- | --- | --- | --- | --- |
| AccD* | AT5G59860 | NA | RNA-binding (RRM/RBD/RNP motifs) family protein | 1.29 | 0.94 | 6.07E-04 | 6.51E-01 | 7.41E-06 |
| AccD* | AT1G16750 | NA | Protein of unknown function, DUF547 | 2.18 | 0.94 | 1.65E-03 | 6.91E-01 | 3.32E-03 |
| Ycf1/2 | AT1G04110 | SDD1 | Subtilase family protein | 0.91 | 0.94 | 1.71E-03 | 3.63E-01 | 1.66E-02 |
| Ycf1/2** | AT5G22450 | NA | unknown protein | 1.68 | 0.81 | 8.14E-03 | 1.92E-01 | 6.25E-02 |
| RNA pol.* | OS03G58204 | NA | NA | 0.39 | 0.75 | 8.50E-03 | 1.97E-01 | 6.91E-03 |
| Ribo. | AT4G14100 | NA | transferases, transferring glycosyl groups | 0.41 | 0.67 | 1.02E-02 | 7.52E-01 | 4.74E-01 |
| RNA pol. | AT3G26618 | ERF1-3 | eukaryotic release factor 1-3 | 0.54 | 0.68 | 1.08E-02 | 7.37E-01 | 6.40E-01 |
| ClpP1** | AT1G09800 | NA | Pseudouridine synthase family protein | 5.28 | 0.74 | 1.23E-02 | 5.69E-01 | 1.53E-03 |
| Ycf1/2 | AT4G25320 | NA | AT hook motif DNA-binding family protein | 0.75 | 0.75 | 2.22E-02 | 2.66E-01 | 2.61E-02 |
| Ycf1/2 | OS03G53360 | NA | NA | 1.59 | 0.88 | 2.22E-02 | 2.19E-01 | 2.14E-01 |
| AccD | AT5G36000 | NA | BEST A.thaliana match: reduced male fertility | 0.18 | 0.80 | 2.65E-02 | 5.53E-01 | 8.69E-03 |
| Ycf1/2* | AT1G55870 | ATPARN, AHG2 | Polynucleotidyl transferase, ribonuclease H-like superfamily protein | 2.61 | 0.73 | 2.85E-02 | 3.37E-01 | 4.26E-03 |
| ClpP1** | AT2G16770 | bZIP23 | Basic-leucine zipper (bZIP) transcription factor family protein | 4.48 | 0.63 | 3.05E-02 | 7.41E-01 | 9.04E-04 |
| RNA pol., AccD, Ribo. | AT4G19985 | NA | Acyl-CoA N-acyltransferases (NAT) superfamily protein | 0.42 | 0.66 | 3.20E-02 | 4.10E-01 | 7.14E-01 |
| RNA pol. | AT1G69410 | ATELF5A-3, ELF5A-3 | eukaryotic elongation factor 5A-3 | 0.12 | 0.61 | 3.20E-02 | 8.76E-01 | 4.79E-01 |
| RNA pol. | AT3G17880 | HIP, ATTDX, ATHIP2, TDX | tetratricopeptide domain-containing thioredoxin | 0.11 | 0.62 | 3.37E-02 | 7.47E-01 | 8.60E-02 |
| AccD | AT4G19350 | EMB3006 | embryo defective 3006 | 1.88 | 0.75 | 3.42E-02 | 7.16E-01 | 1.97E-02 |
| Ycf1/2** | OS03G26080 | NA | NA | 4.10 | 0.71 | 3.50E-02 | 2.20E-01 | 1.17E-01 |
| RNA pol. | AT5G25840 | NA | Protein of unknown function (DUF1677) | 0.45 | 0.63 | 3.54E-02 | 2.60E-01 | 1.98E-01 |
| RNA pol. | AT4G39920 | POR, TFCC | C-CAP/cofactor C-like domain-containing protein | 0.36 | 0.65 | 3.71E-02 | 5.08E-01 | 2.94E-01 |
| Ribo., RNA pol. | AT1G71000 | NA | Chaperone DnaJ-domain superfamily protein | 0.62 | 0.62 | 3.93E-02 | 5.96E-01 | 5.68E-01 |
| AccD | AT5G39420 | cdc2cAt | CDC2C | 4.59 | 0.74 | 3.95E-02 | 6.58E-01 | 4.11E-01 |
| RNA pol. | AT1G03330 | NA | Small nuclear ribonucleoprotein family protein | 0.69 | 0.77 | 4.27E-02 | 2.18E-01 | 1.83E-01 |
| Ribo. | AT2G03820 | NA | nonsense-mediated mRNA decay NMD3 family protein | 0.19 | 0.59 | 4.34E-02 | 7.52E-01 | 1.06E-01 |
| RNA pol. | AT5G26610 | NA | D111/G-patch domain-containing protein | 0.65 | 0.58 | 4.49E-02 | 5.07E-01 | 5.88E-01 |
| Phot., RNA pol. | AT5G20040 | IPT9 | isopentenyltransferase 9 | 0.35 | 0.63 | 4.74E-02 | 1.70E-01 | 1.61E-02 |
| AccD | AT5G52860 | ABCG8 | ABC-2 type transporter family protein | 7.03 | 0.57 | 1.16E-01 | 2.18E-13 | 6.62E-02 |
| AccD | AT2G28315 | NA | Nucleotide/sugar transporter family protein | 6.45 | 0.63 | 1.17E-01 | 2.18E-13 | 1.53E-01 |
| AccD | OS09G39370 | NA | NA | 1.68 | 0.59 | 2.70E-01 | 2.18E-13 | 1.51E-01 |
| MatK | AT4G23330 | NA | BEST A. thaliana match: eukaryotic translation initiation factor 3A | 0.37 | 0.43 | 3.96E-01 | 2.91E-13 | 5.69E-01 |

## Discussion

### Genomic signatures of plastid-nuclear interactions can be detected with ERC in plants

ERC has revealed novel interactions in animals and fungi but, until now, has not been applied at broad phylogenetic scales in plants due to the prevalence of gene/genome duplication. We adapted existing techniques, initially developed with the stringent requirement of one-to-one orthology, to make them more tolerant of duplications, thus allowing us to analyze a substantial portion of plant nuclear genomes. Our pipeline (Fig. 2) included several features tailored to analyze plant genomes. For example, our orthologous subtree extraction procedure identified subtrees with reduced paralogy compared to input trees, shifting the distribution of trees closer to one-to-one orthologous relationships without substantial loss of data (Fig. S1). In addition, our iterative gene tree/species tree (GT/ST) reconciliation approach resolved topological disagreements when they lacked phylogenetic support, allowing us to minimize phylogenetic noise while retaining well-supported phylogenetic signature. The typical implementations of ERC assume every gene tree has the exact same sampling and topology (Nathan L Clark et al., 2012; Findlay et al., 2014; Wolfe & Clark, 2015). However, this is rarely the case in plant datasets, which are prone to topological variation introduced by internal duplications, incomplete lineage sorting, and differential gene loss (Degnan & Rosenberg 2009; Leebens-Mack, Barker, Carpenter et al., 2019), making it infeasible to compare individual branches in a one-to-one fashion between gene trees. This challenge prompted us to apply a root-to-tip approach to calculating branch lengths. A drawback of this approach is that it introduces pseudoreplication via sampling shared internal branches multiple times (Felsenstein, 1985; Yan et al., 2019). We minimized this effect with our taxon-sampling by avoiding closely related species and, thus, approximating a ‘star-phylogeny’ as closely as possible. Finally, when multiple paralogs were present in a gene tree, we averaged the branch lengths between all paralogs for a given species. This approach allowed us to accommodate localized duplication events within trees. Our results offer proof-of-principle that ERC can be successfully extended to plant genomes at phylogenetic scales spanning angiosperm diversity and likely further. While we focused on plastid-nuclear interactions, our results open the door to applying this method broadly to probe the entire plant interactome.

The predominance of known N-pt proteins among our ERC hits validates the use of ERC in detecting functional relationships and illustrates the impacts of plastid-nuclear interactions on evolutionary rates at a genome-wide scale. However, it is important to consider the correlative nature of ERC and the fact that detected effects do not always imply direct functional interactions. For example, we observe significant enrichment of N-mt proteins among our ERC hits (albeit a much weaker signal than for N-pt genes; Fig. 4 and Table 1). Given that our ERC searches were seeded with plastome partitions, it is tempting to interpret these signals as evidence for cofunctionality or crosstalk between mitochondria and plastids. Although such factors may contribute to the observed N-mt signal, the rates of evolution of the plastome and mitochondrial genome are known to be partially correlated with each other. Lineages such as *Plantago, Silene*, and Geraniaceae that exhibit rapid rates of plastome evolution in our sample (Fig. 1) also have unusually rapidly evolving mitochondrial genomes (Cho et al., 2004; Parkinson et al., 2005; Jansen et al., 2007; Mower et al., 2007; Sloan et al., 2009; Seongjun Park et al., 2017). As such, we would expect overlap between ERC hits from the two genomes even in the absence of co-functionality between the mitochondria and plastids. Similarly, our plastome partitions do not evolve entirely independently of each other. Although the magnitudes of rate acceleration can vary greatly among genes (Fig. 1; (Guisinger et al., 2008; Sloan, Triant, Forrester, et al., 2014; Seongjun Park et al., 2017; Shrestha et al., 2019)), we observe significant ERC between all pairs of our plastome partition trees (Table S3), limiting our ability to distinguish specific signatures of ERC for individual partitions. Consistent with this, we found overlap between the hits identified for each partition (Fig. S2A-B). Multiple regression analyses provided some assistance in identifying the partitions making the strongest contributions to plastid-nuclear ERC (Fig. S2C-D, Tables 1 and 2), but further investigation will be needed to tease apart the effects of correlated rates of evolution within and between cytoplasmic genomes in order to pinpoint the loci responsible for ERC with nuclear genes.

### Networks of cofunctional proteins are connected via their involvement in plastid proteostasis

ERC analyses point to plastid proteases, ribosomal proteins (subunits and binding/maturation factors), translation initiation/elongation factors, and proteins involved in protein import into the plastids (Fig. 4, Table 1), all of which contribute to maintenance of protein quality control, proteostasis, and the unfolded protein response (Kim et al., 2013; Dogra et al., 2019) (Fig. S4). Proteases exhibit some of the most striking signatures of ERC. In addition to Clp subunits, we observed strong ERC for FtsH7, FtsH9 and FtsH11. These proteins are thought to form two separate protease complexes, both of which localize to the plastid envelope (Ferro et al., 2003, 2010; Wagner et al., 2012). Interaction partners and substrates have been identified for FtsH11 (Adam et al., 2019), but very little is known about the function of the FtsH7/9 complex. These FtsH protease subunits do not appear to form a complex with any plastid-encoded protein, making them an example of correlated plastid-nuclear evolution in the absence of direct physical interaction. It is somewhat surprising that we did not observe significant ERC for other members of the gene family that comprise the thylakoid FtsH protease (FtsH1/2/5/8) considering that Clp mutants are suppressors of variegation phenotypes in thylakoid FtsH mutants (Sungsoon Park & Rodermel, 2004; Yu et al., 2008). However, our results may be consistent with the prior observation that expression of thylakoid FtsH subunits are unaffected by Clp mutants, suggesting a lack of reciprocity in the interactions between Clp and the thylakoid FtsH protease (Kim et al., 2013). On the other hand, we do observe strong ERC for additional members of the FtsH family, FtsH12 and FtsHi5, which form part of a complex that facilitates protein import across the inner membrane of the plastid, acting as an ATPase motor rather than a protease (Kikuchi et al., 2018). Plastid-nuclear ERC for this complex may result from the fact that it also contains plastid-encoded Ycf2 (another FtsH paralog) (Kikuchi et al., 2018). These and other genes involved in protein import (most notably, TIC110) (Table 1) point to the strong signature of plastid-nuclear evolution exhibited by import machinery, again highlighting the prominence of proteostasis pathways in our ERC hits.

We observed ERC for several plastid ribosomal subunits and other genes involved in plastid translation (Table 1). For example, SVR7 is a pentatricopeptide repeat (PPR) protein that is involved in plastid rRNA processing, which (like Clp subunits) suppresses thylakoid FtsH mutant variegation (Liu et al., 2010), again pointing to functional connections between plastid translation and other proteostasis pathways. However, perhaps our most surprising piece of evidence for the role of translation in plastid-nuclear ERC is the association between ClpP1 and protein subunits of the cytosolic ribosome (Fig. 4 and Fig. S3). While ERC has been previously detected among cytonuclear subunits in plastid and mitochondrial ribosomes (Sloan, Triant, Wu, et al., 2014; Weng et al., 2016), the cytosolic ribosomes themselves have never been demonstrated to exhibit ERC with the mitogenome or plastome. Most of the plastid proteome is synthesized in the cytosol, meaning the levels of N-pt and plastid-encoded proteins must be regulated to achieve stoichiometric balance for cytonuclear complexes (Colombo et al., 2016). In mitochondria, this balance is achieved through coordination of cytosolic and mitochondrial translation (Houtkooper et al., 2013; Couvillion et al., 2016). Recent evidence suggests that changes in cytosolic translation may have strong genetic interactions with plastid proteostasis machinery. Specifically, mutation of a cytosolic ribosome subunit was shown to enhance variegation phenotypes in thylakoid FtsH mutants (Wang et al., 2018). Given that disruption of plastid translation can suppress these same phenotypes (Yu et al., 2008; Liu et al., 2010; Zheng et al., 2016), it appears that ribosomes in both compartments play a key role in maintenance of plastid-nuclear stoichiometric balance. Additionally, we observe strong ERC for a putative tRNA pseudouridine synthase (AT1G09800) that shows no evidence of plastid or mitochondrial targeting (Table 2), meaning it likely modifies cytosolic tRNAs, again consistent with cytosolic translation being subject to plastid-nuclear selection. These results suggest that the effects of perturbation in plastid proteostasis may extend to cytosolic ribosomes, supporting a level of cofunction-mediated ERC that spans cellular compartments.

Genes involved in various aspects of proteostasis appear to have been subject to accelerated protein evolution in independent angiosperm lineages. We propose that proteostasis systems have been perturbed in these lineages, causing shifts in selection that simultaneously affected numerous functionally related genes. Although the evolutionary events that may have led to these changes are unclear, one possible explanation could be related to the constant stoichiometric pressure plants experience in the face of nuclear gene/genome duplication (Birchler & Veitia, 2012; Sharbrough et al., 2017). Similarly, the susceptibility of plastomes to instability and rearrangements in certain angiosperm lineages (Jansen et al., 2007) could provide an initial trigger that elicits a series of coevolutionary responses. It has also been hypothesized that antagonistic interactions between the nucleus and selfish genetic elements in the plastids could drive accelerated rates of evolution (Rockenbach et al., 2016; Sobanski et al., 2019). Finally, perturbations could be prompted by changes in abiotic or biotic stress, as many of the pathways that contribute to proteostasis are stress-responsive (e.g., the unfolded protein response to photooxidative stress) (Dogra et al., 2019). The cause of these perturbations may differ by lineage and disentangling them could reveal a critical driver of plant genome evolution. Regardless of the mechanisms, it is striking that the ripple effects are apparent across disparate pathways and cellular compartments and can be detected against the background of the entire genome in a large swath of plant diversity.

### ERC points to novel plastid-nuclear interactions

Decades of proteomics research have led to the identification of more than 2,400 plastid-localized proteins in *Arabidopsis* (http://ppdb.tc.cornell.edu; http://cymira.colostate.edu/). Yet, these proteins may only represent about 70% of the plastid proteome (Millar et al., 2006; van Wijk & Baginsky, 2011; Christian et al., 2020). Large-scale plastid proteomic surveys are limited by ascertainment bias associated with protein expression level, tissue- and condition-specificity of expression/plastid-localization, and biochemical properties that impact mass spectrometry (van Wijk & Baginsky, 2011). ERC offers an alternative line of evidence for plastid function/localization that is complementary to biochemical approaches and may not share the same biases. Our analyses returned several proteins that lack plastid-targeting annotations (Table 2) and represent candidates for novel N-pt proteins. For example, two of our strongest non-plastid-localized hits are annotated as RNA-binding (AT5G59860) and GPI-anchored adhesin-like (AT1G16750) proteins based on *in silico* predicted domains but are, otherwise, lacking in functional information. The signature of plastid-nuclear ERC that we observe for the genes in Table 2 suggests they have experienced correlated changes selection associated with accelerated plastome evolution. A natural hypothesis is that these are cryptic N-pt proteins that have evaded biochemical identification. However, an alternative explanation is that they contribute to plastid function without localizing to plastids, similar to our hypothesis for cytosolic ribosomes and the pseudouridine synthase described above. A third possibility is that the proteins are plastid-localized in many plants but not in *Arabidopsis*, which is possible given the apparent lability of plastid-targeting across plants (Christian et al., 2020; Costello et al., 2020). While each of these explanations come with their own functional and evolutionary implications, future work to disentangle these alternative hypotheses will undoubtably advance our understanding of the full repertoire of plastid-nuclear interactions.

## Methods

### Taxon sampling and obtaining sequence data

Our analysis was conducted on publicly available genomes and transcriptomes. Because the signature of ERC relies on phylogenetic rate heterogeneity, we sampled species that are known to exhibit differences in evolutionary rate for at least some plastid genes, including representatives from accelerated lineages, such as *Acacia aulacocarpa, Oenothera biennis, Geranium maderense, Plantago maritima, Lobelia siphilitica, Silene noctiflora*, and *Oryza sativa* (Jansen et al., 2007; Guisinger et al., 2008; Knox, 2014; Sloan, Triant, Forrester, et al., 2014; Dugas et al., 2015; Nevill et al., 2019; Shrestha et al., 2019). We also sampled species that exhibit the slow background rate of plastome evolution typical for most angiosperms. We did not include parasitic species with accelerated plastome evolution, as these represent special cases of plastid evolution associated with loss of photosynthetic function (Wicke et al., 2016).

Because our ERC analysis employs a root-to-tip strategy for measuring branch lengths (described below), we avoided sampling pairs of species that are closely related to each other in order to minimize pseudoreplication caused by shared internal branches (Felsenstein, 1985; Yan et al., 2019). In addition to the 18 species chosen by the above logic, we included *Amborella trichopoda* and *Liriodendron chinense* as outgroups. We chose to include two outgroups so gene families would contain an outgroup sequence even if gene loss occurred in one of the two species, allowing us to analyze a larger proportion of gene families. It should be noted that phylogenetic placement of magnoliids (including *Liriodendron*) with regard to the ingroup (eudicots and monocots) has been a topic of debate (Soltis et al., 1999; Zanis et al., 2002; Hilu et al., 2003; Qiu et al., 2005, 2006). However, large-scale analysis of the plastid genome resolved *Liriodendron* as an outgroup to a eudicot/monocot clade (Jansen et al., 2007).

We obtained the full set of twenty proteomes from several sources (Table S4) and processed fasta files to add standardized sequence identifiers. For genome-based datasets that contained multiple splice variants per gene, we used only the first gene model (i.e. gene model ending in .1) and removed the rest to avoid falsely defining splice variants as paralogs in gene family clustering.

Plastome gene datasets were extracted from GenBank files (see Table S4) using a custom BioPerl script and manually curated to deal with missing annotations and inconsistent naming conventions. The corresponding protein sequences were either analyzed individually (ClpP1, AccD, and MatK) or concatenated from multiple plastid genes that are part of a common plastid complex and/or pathway (photosynthesis, ribosomes, RNA polymerase, and Ycf1/Ycf2) (Table S1). The plastome sampling matched the nuclear proteome samples described above except that no plastome sequence was available for *Acacia aulacocarpa*, so we used the *Acacia ligulata* plastome in its place. The *accD* gene is missing from the plastome of *Oryza sativa* and *Lobelia siphilitica*, and *ycf1* and *ycf2* are missing from *Oryza sativa* and *Geranium maderense.* These species were omitted from the alignments and trees for AccD and Ycf1/Ycf2. Amino acid alignments based on plastome partitions were used to estimate branch lengths on a constraint tree with a topology based on Angiosperm Phylogeny Website (http://www.mobot.org/MOBOT/research/APweb) (Fig. 1).

### Gene family clustering, sequence alignment, and phylogenetic inference

We clustered homologous gene families using Orthofinder (v2.2.6) (Emms & Kelly, 2015) and performed multiple sequence alignment using the L-INS-i algorithm in MAFFT (v7.407) (Katoh & Standley, 2013). We used RAxML (v8.2.12) (Stamatakis, 2014) to infer maximum likelihood trees with 100 bootstrap replicates. Tree inference was performed using the command below for each gene. The -m argument indicates the model used (gamma distributed rate heterogeneity, empirical amino-acid frequencies, and the LG substitution model). The -p argument provides a seed for parsimony search. The -x argument provides a seed for rapid bootstrapping. The -# argument indicates the number of bootstrap replicates. The -f a argument implements rapid bootstrap analyses and best scoring tree search. The -T argument indicates the number of threads used for parallel computing.

raxmlHPC-PTHREADS-SSE3 -s <input file name> -n <output file name> -m PROTGAMMALGF -p 12345 -x 12345 -# 100 -f a -T 24

For the step in which we optimized branch lengths on a constraint tree (see below), we used the following command, with -f e indicating parameter and branch-length optimization.

raxmlHPC-PTHREADS-SSE3 -s <input file name> -n <output file name> -t <name of constraint tree file> -m PROTGAMMALGF -p 12345 -T 24 -f e

### Subtree extraction and quality control pipeline

ERC analyses are sensitive to false inferences of orthology. Particularly, treating cryptic out-paralogs as orthologs can alter branch length estimates (Smith & Hahn, 2020). While Orthofinder clusters sequences that share homology, these clusters do not always represent groups that share strict orthology. ERC analyses are also sensitive in poorly aligned sequences, which can result in long outlier branches on trees. To address these inherent challenges to genome-scale phylogenetic analyses, we built a pipeline to process nuclear gene trees and retain the portions of alignments and trees least likely to be affected by biasing factors. Our pipeline enlists several existing programs. In this section we provide a summary of the steps in the pipeline and point the reader to subsequent sections for details on our application of individual components of the pipeline.

**(Step 1)** Starting with the full gene trees we performed GT/ST reconciliation in order to root the tree, rearrange poorly supported portions of the tree to conform with the species tree, and infer nodes in the tree that represent gene duplication rather than speciation. **(Step 2)** We used duplication information from step 1 to extract subtrees representing orthology groups. **(Step 3)** We performed a second round of sequence alignment (using MAFTT as above) to generate alignments that contain only the sequences in subtrees. **(Step 4)** We trimmed these alignments to remove poorly aligned regions using GBLOCKS. We filtered out any alignments with a length of less that 50 amino acids as well any alignments for which GBLOCKS trimming resulted in the removal of an entire sequence from the alignment. **(Step 5)** We inferred a new phylogeny for each subtree from the trimmed alignment using RAxML as above and again applied GT/ST reconciliation to the subtree trees to rearrange poorly supported nodes and root the tree. **(Step 6)** We used the reconciled versions of the gene trees (as constraint trees) and the trimmed version of the alignments to optimize final branch lengths for use in downstream ERC analyses. **(Step 7)** As a final means of quality control before performing ERC analyses, we assessed each tree to ask whether the ingroup forms a monophyletic clade in the branch-length-optimized tree. Those that were not monophyletic were pruned and rerooted in order to retain ingroup monophyly. We also filtered out trees with one very long outlier branch by removing any trees in which the longest branch is more than ten times the length of the second longest branch.

### GT/ST reconciliation

We used GT/ST reconciliation to reconstruct the history of gene duplication for each gene tree using Notung (v2.9) (Vernot et al., 2008; Stolzer et al., 2012). Briefly, Notung compares the topology of a gene tree inferred from an individual gene to the topology of a user-input species tree. We used the topology of the plastome trees described above as our species tree. Incongruencies between the gene tree and species tree are taken to be the result of historical gene duplication occurring at specific nodes of the tree. Notung uses a parsimony framework to reconcile these incongruences by inferring duplication and loss events along the gene tree to yield the most parsimonious series of duplication and loss events for each gene tree. Notung can also apply this logic to root unrooted gene trees by the most parsimonious root. Since topological incongruence is the signature by which Notung infers duplication events, inferences are sensitive to phylogenetic error, evidenced by branches with low bootstrap support. To avoid false inference of duplication from weakly supported branches, we made use of Notung’s option to only infer duplication that is supported by branches with bootstrap support of at least 80 percent.

We performed the rearranging step for each gene tree on the command line with the following command:

java -jar Notung-2.9.jar <path to gene tree file> -s <path to species tree file> --rearrange -- threshold 80 --treeoutput nhx --nolosses --speciestag prefix --edgeweights name --outputdir <output directory>

We performed the rooting step for each gene tree with the following command:

java -jar Notung-2.9.jar <path to rearranged gene tree file> -s <path to species tree file> --root -- treeoutput nhx --nolosses --speciestag prefix --edgeweights name --outputdir <output directory>

In both of the above commands, --treeoutput nhx indicates trees to be output in the newick extended format, which allows for the retention of duplication information. --nolosses indicates that loss information is omitted from the output file (but still included in the reconciliation process). --speciestag and --edgeweights instructs Notung where to find relevant information in the input file.

### Orthologous subtree extraction

We used duplication information from Notung to extract portions of gene trees (i.e. subtrees) in which the taxa share orthology relationships to each other (as opposed to paralogy). We required that these subtrees contain at least one eudicot, one monocot, and one outgroup sequence (*Amborella trichopoda* or *Liriodendron chinense*). We required that at least ten species be represented in each subtree and the eudicot and monocot taxa in the subtree (i.e. the ingroup) form a monophyletic clade. To extract subtrees that fulfill these criteria, for each gene tree we started by iteratively splitting the tree at each node indicated as a duplication node by Notung and retaining the two daughter trees from the splits. Daughter trees were assessed independently and those that fulfilled the above criteria were retained, meaning that multiple subtrees were retained from an initial gene tree in some cases. The final subtrees retained after this process were non-overlapping subtrees containing at least ten taxa representing eudicots, monocots, and at least one outgroup with eudicots and monocots forming a monophyletic clade.

### Multiple sequence alignment trimming with GBLOCKS

We used GBLOCKS (v0.91b) (Castresana, 2000) to trim poorly aligned regions of our alignments using the below command, with -b4 indicating the minimum length of the retained block, -b5=h indicating that gaps are allowed in up to half of the total species, and -b2 indicating the minimum number of sequences for a flank position.

Gblocks <aln directory> <aln file name> -b5=h -b4=5 -b2=<half the total number of sequences>

### Rerooting to retain ingroup monophyly following subtree phylogenetic inference

We realigned and inferred a new phylogeny for subtrees using the same methodology described above. In some cases, these new trees no longer placed eudicots and monocots (i.e. the ingroup) as a monophyletic group, which is a requirement of our downstream ERC analyses. This problem arose in trees in which there were multiple sequences from outgroup species and one or more of these taxa was nested within the ingroup causing the ingroup to be polyphyletic. For these trees, we identified the offending outgroup branches and pruned them from the tree. If *Amborella trichopoda* remained following pruning, we rooted on a branch leading to that species, choosing one at random if there were multiple *Amborella trichopoda* sequences. If no *Amborella trichopoda* branches remained, we rooted on *Liriodendron chinense* in a similar fashion.

### ERC analysis

Branch lengths for ERC analyses were obtained from rooted branch-length-optimized gene trees. We used a root-to-tip method that measures the collective lengths of the path of branches from each ingroup tip to the node representing the most recent common ancestor of all ingroup tips. We obtained root-to-tip branch length measurements for all ingroup species for each gene tree using *dist.nodes()* command from the *Ape* package (Paradis et al., 2004) in *R*. When multiple paralogs from a given species were present, the mean root-to-tip distance from all paralogs was used. When species were absent from trees, branch lengths were indicated as missing values for those species and excluded from ERC analysis for those genes. To account for lineage-specific differences in whole genome rate of evolution, we normalized the branch length for each species be dividing the value for each tree by the average branch length for that species across all genes in our analysis. These normalized branch length values were used for pairwise ERC comparisons.

We compared each of the seven plastome partition trees against all nuclear trees. Each pairwise comparison comprised a correlation analysis of the branch lengths for each species in the plastid tree versus the branch lengths for the same species in the nuclear gene tree (see Fig. 3 for visual depiction). For each pairwise comparison we calculated Pearson and Spearman correlation coefficients. Because there is no clear biological expectation for significant inverse relationships in ERC, we only considered genes with positive correlations (slope > 0) in downstream analyses. We adjusted p-values for multiple comparisons using the false discovery rate (FDR) method implemented with the *p.adjust()* function in *R*.

### CyMIRA and Gene Ontology functional enrichment analyses

In order to perform functional enrichment analyses, we needed a threshold to separate our ‘hits’ from our background genes. We chose to make use of p-values from both Pearson correlation and Spearman correlation as metrics because Pearson gains power from large branch lengths, potentially expected under true evolutionary co-acceleration, and Spearman is less sensitive to outlier branches. Any gene with a Pearson p-value ≤ 0.05 and a Spearman p-value ≤ 0.1 was designated as a ‘hit’. Our goal here was to identify the tail of the distribution for the sake of functional enrichment analysis. A more stringent threshold was applied when assessing the significance of individual hits (Table 1 and 2).

We used the *Arabidopsis* sequence identifiers present within gene families to probe functional enrichment of significant hits based on localization/interaction annotations from *CyMIRA* and functional annotations from Gene Ontology. We used the 7929 genes in our filtered dataset as the background (rather than using the full *Arabidopsis* genome). For gene families that contained multiple *Arabidopsis* paralogs, we selected a single *Arabidopsis* paralog at random to represent the family. Families that did not contain any *Arabidopsis* sequences were omitted from this portion of the analysis. Fold enrichment was calculated as number of observed hits in a category divided by the number of expected hits in a category, where the expected is the proportion of the background in a category multiplied by the number of hits. The localization/interaction enrichment analyses were performed in *R*. Gene Ontology enrichment analyses was performed using the *PANTHER* web-based tool (http://geneontology.org/) (database release from 10-08- 2019). Significance of enrichment was assessed with Fisher’s Exact Test with an FDR correction for multiple comparisons.

### Identification of genes displaying strong signatures of ERC

To identify individual genes displaying the strongest signatures of plastid-nuclear ERC, we applied more stringent criteria that considered Pearson and Spearman correlation p-values in their raw and FDR-corrected forms. Our criteria for labeling a gene as a strong hit is that either the adjusted Pearson p-value or the adjusted Spearman p-value (or both) must be ≤0.05. Additionally, for the genes in which only one of the two adjusted p-values was ≤0.05, we also required that the raw Pearson and raw Spearman p-value both be ≤0.05. This approach allowed us to incorporate information from both correlation coefficients and from FDR multiple test correction while still retaining power to detect the strongest hits. Genes passing these criteria are presented in Table 1 and 2.

### Multiple regression analyses

To investigate the relative contributions of each plastome partition to the evolutionary rates of each nuclear-encoded protein, we conducted a multiple regression analysis using branch lengths from our constructed trees. Due to the lack of *accD* in *Oryza sativa* and *Lobelia siphilitica* and the lack of *ycf1/ycf2* in *Oryza sativa* and *Geranium maderense*, we excluded branch lengths those three species, which allowed us to include all seven plastome partitions. Each nuclear gene was analyzed separately, where the y values were the normalized branch lengths for each species for that particular gene and the x values were the normalized branch lengths for each plastome partition for each species. Any additional missing data led to removal of the involved species. Models were created using the *lm()* function in *R* with default parameters.

## Author contributions and acknowledgements

E.S.F., A.M.W., and D.B.S. conceived this work and performed analyses. E.S.F. drafted this manuscript with input from all authors. This work was supported by a National Science Foundation (NSF) grant (MCB-1733227) and graduate fellowships from NSF (DGE-1321845) and the National Institutes of Health (T32-GM132057). We thank M.P. Simmons and J.C. Havird for helpful discussion.

## Figures

**Figure S1.**
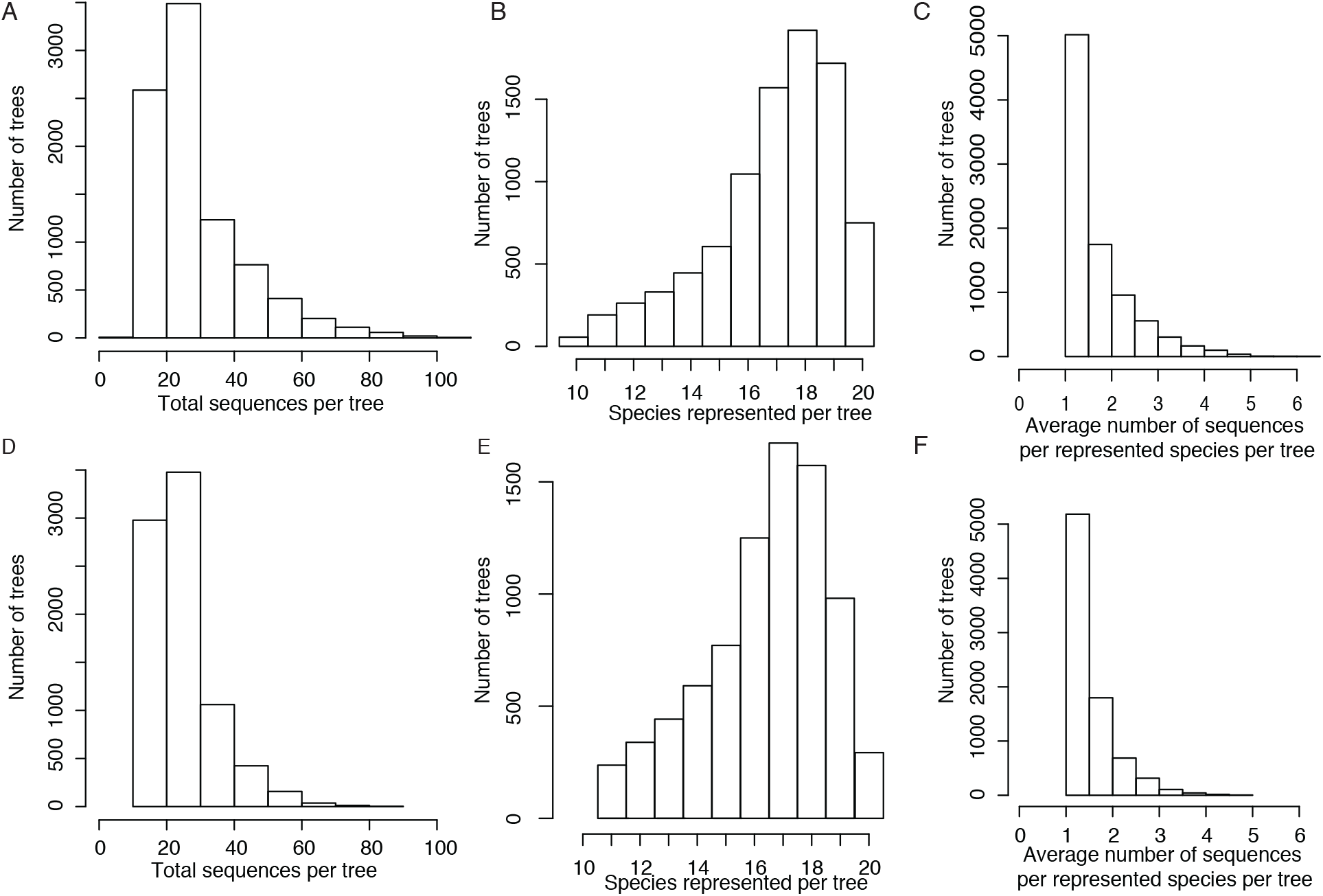
Taxon composition of trees and subtrees. Histograms representing total number of sequences per tree (A and D), number of species represented per tree (B and E), and average number of sequences per represented species per tree (C and F). Distributions are shown for original trees before subtree extraction (see Methods) (A-C) as well as for final subtrees after orthologous subtree extraction (D-F).

**Figure S2.**
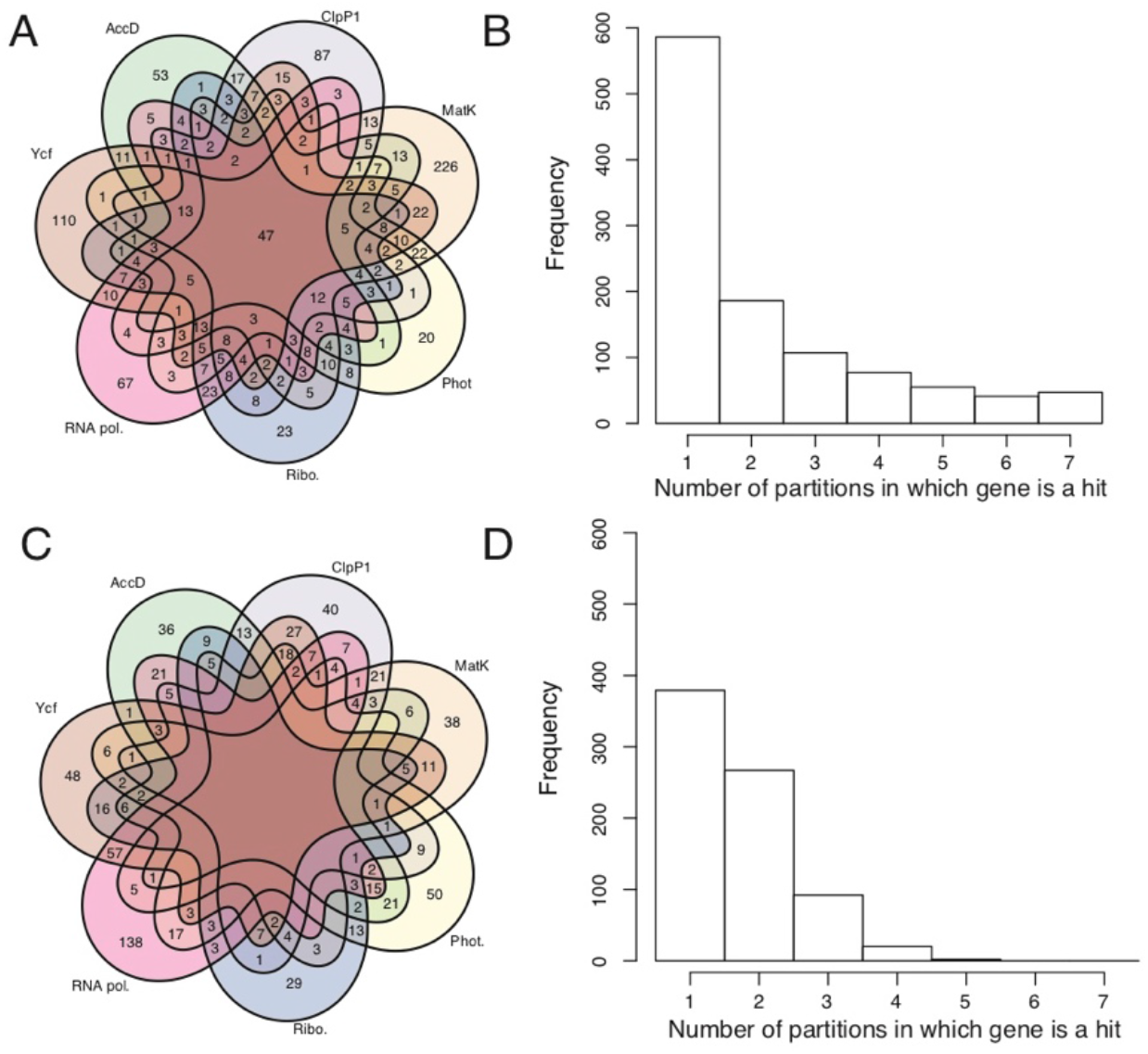
Plastome partition ERC result overlap and multiple regression analysis. (A-B) ERC hit overlap for the single-correlation analyses used in Figs. 4 and 5. (A) Venn diagram showing the overlap of ERC hits between partitions. (B) Histogram showing the number of partitions in which each nuclear gene was a hit. (C-D) ERC hit overlap for the multiple regression analyses. (C) Venn diagram of the overlap of ERC hits between partitions. (D) Histogram of the number of partitions in which each nuclear gene was a hit.

**Figure S3.**
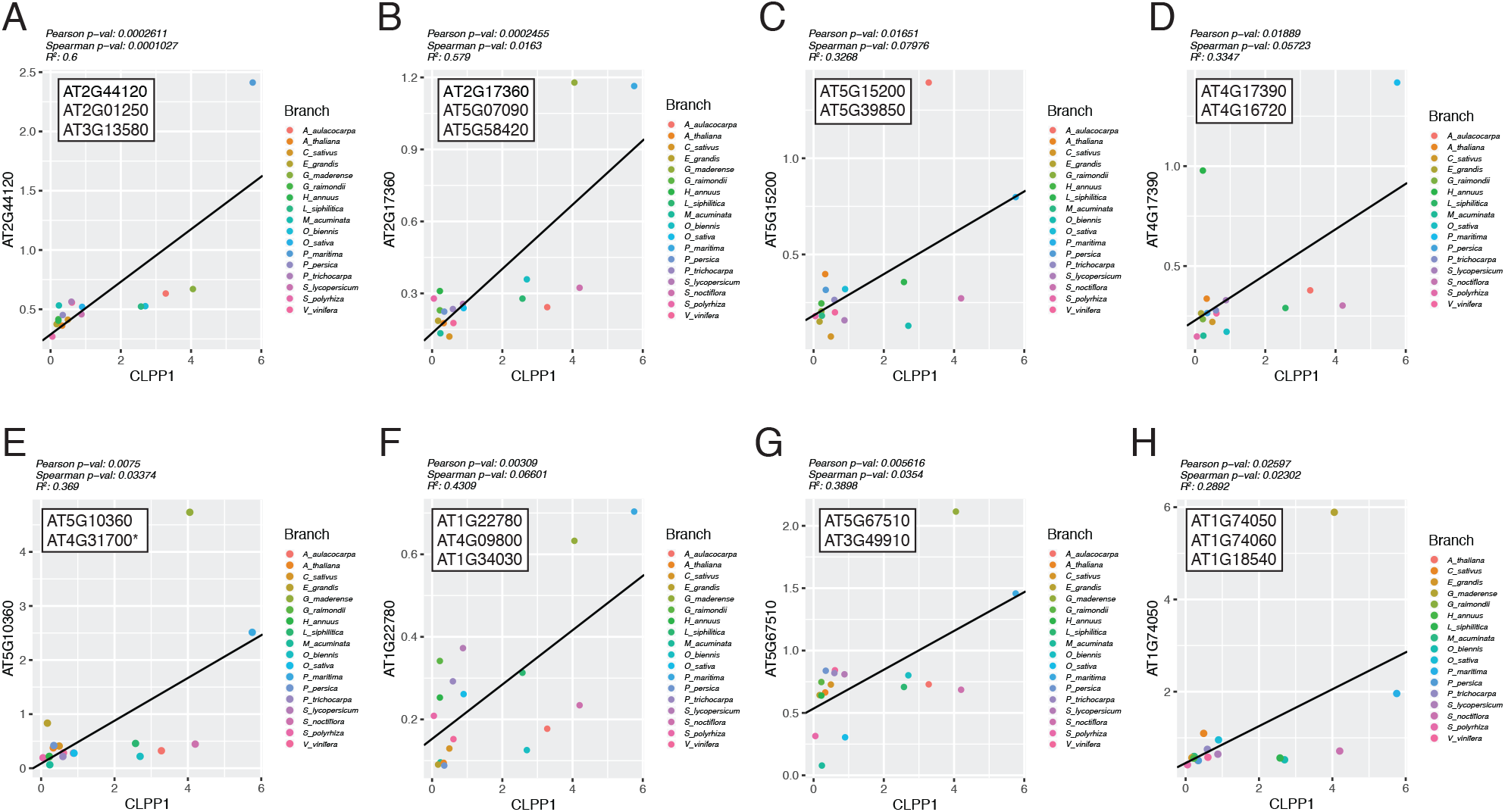
Cytosolic ribosome subunits found to have significant ERC with ClpP1. Correlation plots comparing normalized branch lengths from ClpP1 versus normalized branch lengths for cytosolic ribosome gene trees. All families contained multiple *Arabidopsis* paralogs. The y-axis labels indicate the AGI locus identifier for the randomly chosen paralog used for enrichment analyses. The white box insets list AGI locus identifiers for all Arabidopsis paralogs in each family. All loci shown were compared against previous datasets (Bonen & Calixte, 2005; Tiller et al., 2012; Sloan, Triant, Forrester, et al., 2014; Bieri et al., 2017; Boerema et al., 2018; Waltz et al., 2019) and found to be annotated as cytosolic ribosomes except for AT4G31700 (indicated with *), which was not annotated as a ribosome subunit in the above studies but is annotated at a cytosolic ribosomal subunit elsewhere (Creff et al., 2010).

**Figure S4.**
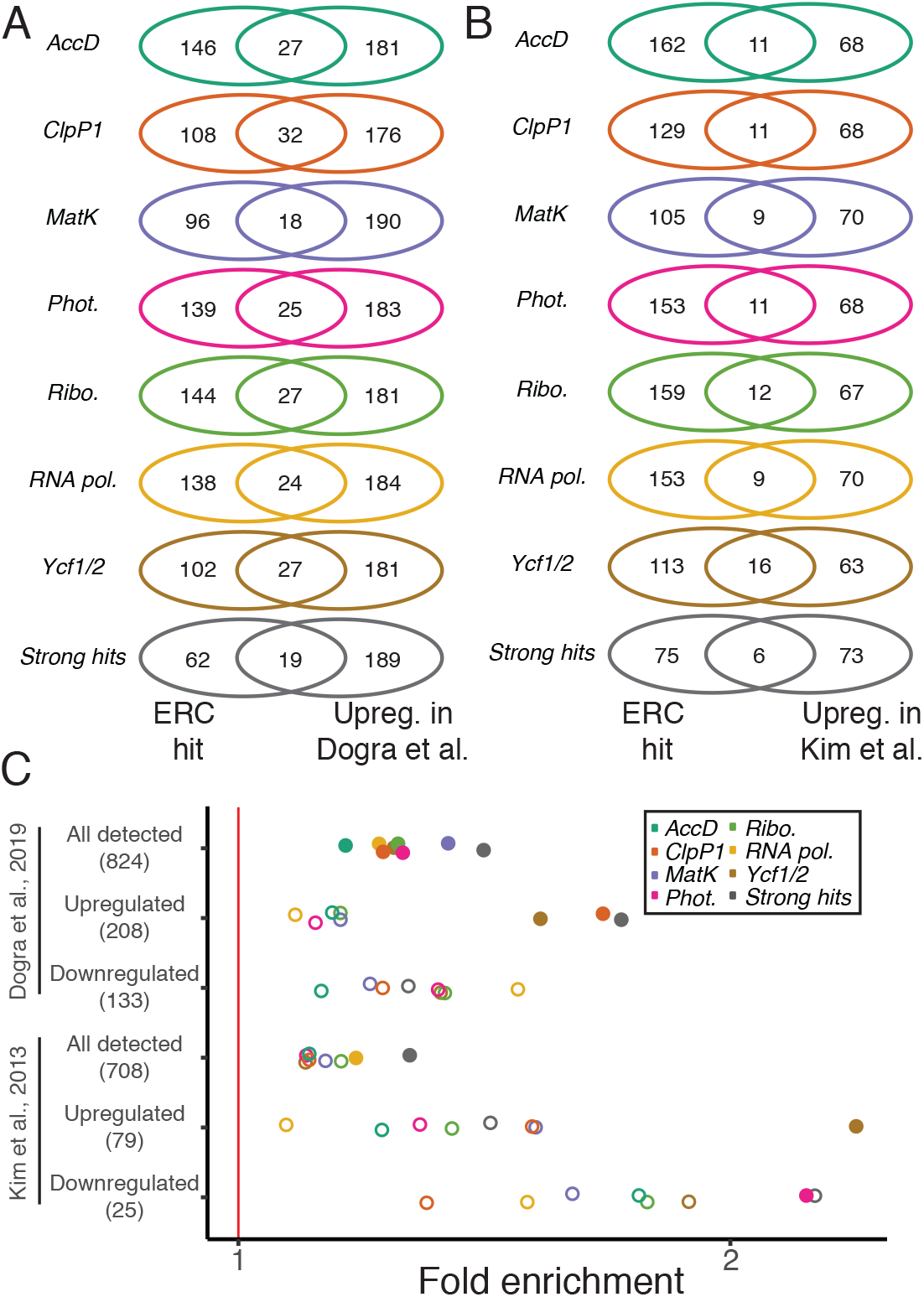
Comparison of ERC hits to genes with altered expression in proteostasis mutants. (A-B) Venn diagrams depicting overlap of ERC hits with upregulated chloroplast proteins from mutants for the metalloprotease subunit FtsH2 (Dogra et al., 2019) (A) and *ClpP3* (Kim et al., 2013) (B). (C) Enrichment analyses testing whether proteins detected with differential expression are enriched among ERC hits. For all panels, ERC hits were filtered to include only chloroplast-localized proteins according to *CyMIRA* (Forsythe et al., 2019).

**Table S1:** Plastome partition multiple sequence alignments. Information about plastome partitions used to infer plastid trees.

| Plastome partition | Number of genes | Gene(s) | Alignment length (AAs) | Missing species |
| --- | --- | --- | --- | --- |
| AccD | 1 | AccD | 1281 | *Oryza sativa* and *Lobelia siphilitica* |
| ClpP1 | 1 | ClpP1 | 177 | NA |
| MatK | 1 | MatK | 458 | NA |
| Photosynthesis | 46 | AtpA, AtpB, AtpE, AtpF, AtpH, AtpI, NdhA, NdhB, NdhC, NdhD, NdhE, NdhF, NdhG, NdhH, NdhI, NdhJ, NdhK, PetA, PetB, PetD, PetG, PetL, PetN, PsaA, PsaB, PsaC, PsaI, PsaJ, PsbA, PsbB, PsbC, PsbD, PsbE, PsbF, PsbH, PsbI, PsbJ, PsbK, PsbL, PsbM, PsbN, PsbT, PsbZ, RbcL, Ycf3, Ycf4 | 10437 | NA |
| Ribosomes | 20 | Rpl14, Rpl16, Rpl2, Rpl20, Rpl22, Rpl32, Rpl33, Rpl36, Rps11, Rps12, Rps14, Rps15, Rps16, Rps18, Rps19, Rps2, Rps3, Rps4, Rps7, Rps8 | 2492 | NA |
| RNA pol. | 4 | RpoA, RpoB, RpoC1, RpoC2 | 3209 | NA |
| Ycf1/Ycf2 | 2 | Ycf1, Ycf2 | 2479 | *Oryza sativa* and *Geranium maderense* |

**Table S2.** ERC hits identified for each plastome partition. Number of hits designated for each partition according to different thresholds referenced throughout the manuscript. *Single correlation* hits were used in Fig. 4 and 5. *Single correlation (strong hits)* were used for Table 1 and 2. *Multiple regression* hits were used in Fig. S2C-F.

|  | **AccD** | **ClpP1** | **MatK** | **Phot.** | **Ribo.** | **RNA pol.** | **Ycf1/2** |
| --- | --- | --- | --- | --- | --- | --- | --- |
| Single correlation | 514 | 371 | 333 | 324 | 322 | 289 | 284 |
| Single correlation (strong hits) | 48 | 21 | 6 | 33 | 59 | 60 | 38 |
| Multiple regression | 159 | 182 | 131 | 160 | 118 | 303 | 226 |

**Table S3.** ERC comparisons among the seven plastome partitions. R^2^ values (top) and Pearson p-values (bottom in parentheses) for the ERC comparisons of plastome partitions to each other. All p-values remained significant (p ≤ 0.05) after FDR correction using the p-values posted in this table.

|  | AccD | ClpP1 | MatK | Phot. | Ribo. | RNA pol. | Ycf1/2 |
| --- | --- | --- | --- | --- | --- | --- | --- |
| AccD | - | - | - | - | - | - | - |
| ClpP1 | 0.58<br>(5.92E-04) | - | - | - | - | - | - |
| MatK | 0.27<br>(3.77E-02) | 0.25<br>(3.41E-02) | - | - | - | - | - |
| Phot. | 0.64<br>(1.94E-04) | 0.37<br>(7.63E-03) | 0.74<br>(4.90E-06) | - | - | - | - |
| Ribo. | 0.72<br>(3.06E-05) | 0.50<br>(9.61E-04) | 0.51<br>(8.50E-04) | 0.83<br>(1.64E-07) | - | - | - |
| RNA pol. | 0.70<br>(5.55E-05) | 0.38<br>(6.29E-03) | 0.41<br>(4.26E-03) | 0.65<br>(5.24E-05) | 0.82<br>(2.55E-07) | - | - |
| Ycf1/2 | 0.79<br>(9.94E-06) | 0.40<br>(9.11E-03) | 0.33<br>(1.91E-02) | 0.68<br>(9.06E-05) | 0.79<br>(3.72E-06) | 0.64<br>(1.85E-04) | - |

**Table S4.**
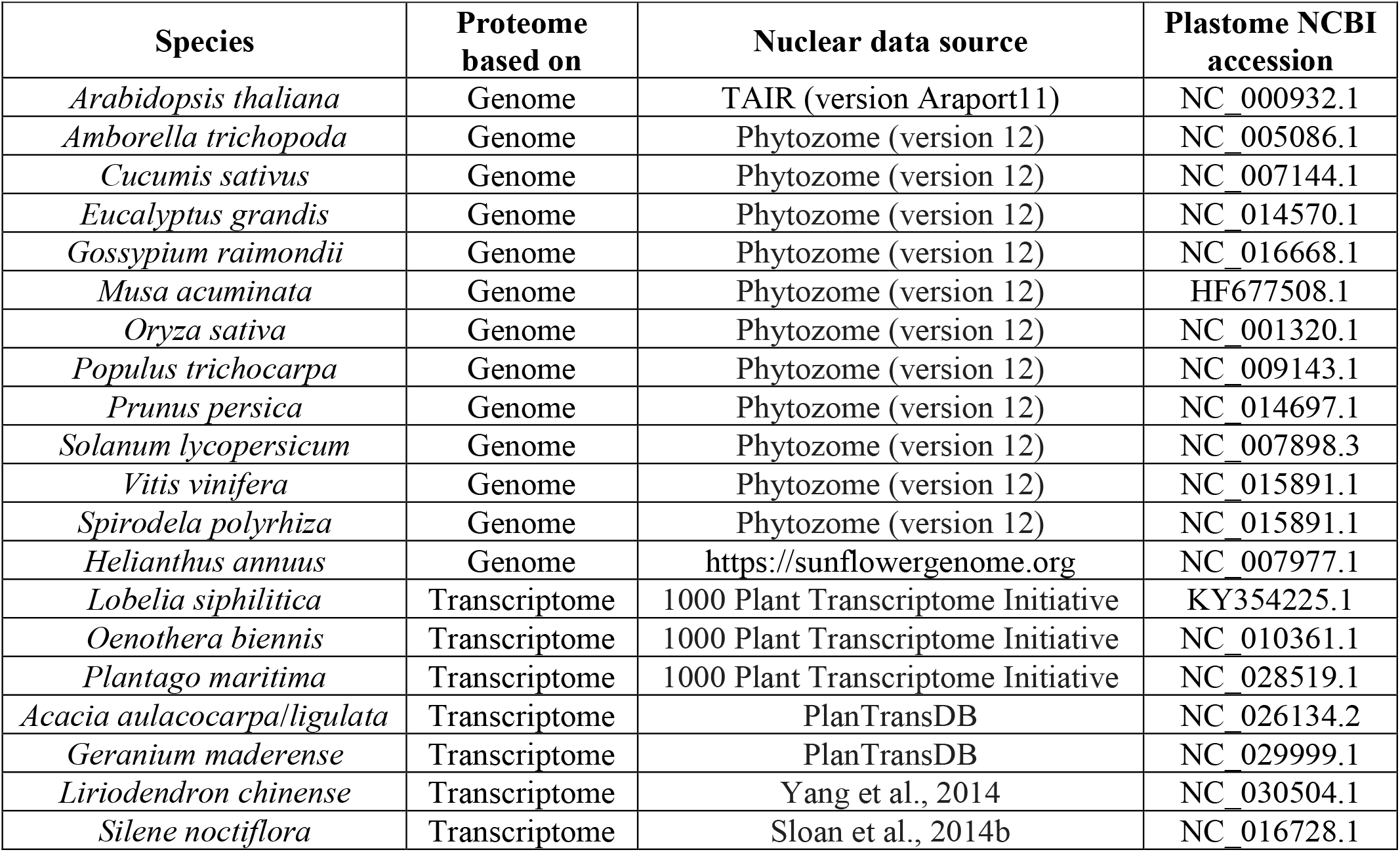
Proteome data sources.

| **Species** | **Proteome based on** | **Nuclear data source** | **Plastome NCBI accession** |
| --- | --- | --- | --- |
| *Arabidopsis thaliana* | Genome | TAIR (version Araport11) | NC 000932.1 |
| *Amborella trichopoda* | Genome | Phytozome (version 12) | NC 005086.1 |
| *Cucumis sativus* | Genome | Phytozome (version 12) | NC 007144.1 |
| *Eucalyptus grandis* | Genome | Phytozome (version 12) | NC 014570.1 |
| *Gossypium raimondii* | Genome | Phytozome (version 12) | NC 016668.1 |
| *Musa acuminata* | Genome | Phytozome (version 12) | HF677508.1 |
| *Oryza sativa* | Genome | Phytozome (version 12) | NC 001320.1 |
| *Populus trichocarpa* | Genome | Phytozome (version 12) | NC 009143.1 |
| *Prunus persica* | Genome | Phytozome (version 12) | NC 014697.1 |
| *Solanum lycopersicum* | Genome | Phytozome (version 12) | NC 007898.3 |
| *Vitis vinifera* | Genome | Phytozome (version 12) | NC 015891.1 |
| *Spirodela polyrhiza* | Genome | Phytozome (version 12) | NC 015891.1 |
| *Helianthus annuus* | Genome | https://sunflowergenome.org | NC 007977.1 |
| *Lobelia siphilitica* | Transcriptome | 1000 Plant Transcriptome Initiative | KY354225.1 |
| *Oenothera biennis* | Transcriptome | 1000 Plant Transcriptome Initiative | NC 010361.1 |
| *Plantago maritima* | Transcriptome | 1000 Plant Transcriptome Initiative | NC 028519.1 |
| *Acacia aulacocarpa/ligulata* | Transcriptome | PlanTransDB | NC 026134.2 |
| *Geranium maderense* | Transcriptome | PlanTransDB | NC 029999.1 |
| *Liriodendron chinense* | Transcriptome | Yang et al., 2014 | NC 030504.1 |
| *Silene noctiflora* | Transcriptome | Sloan et al., 2014b | NC 016728.1 |

## References

Adam, Z., Aviv-Sharon, E., Keren-Paz, A., Naveh, L., Rozenberg, M., Savidor, A., & Chen, J. (2019). The chloroplast envelope protease FTSH11 – Interaction with CPN60 and identification of potential substrates. Frontiers in Plant Science, 10(April), 1–11. https://doi.org/10.3389/fpls.2019.00428

Bansal, M. S., & Eulenstein, O. (2008). The multiple gene duplication problem revisited. Bioinformatics (Oxford, England), 24(13), i132–8. https://doi.org/10.1093/bioinformatics/btn150

Barnard-Kubow, K. B., So, N., & Galloway, L. F. (2016). Cytonuclear incompatibility contributes to the early stages of speciation. Evolution, 70(12), 2752–2766. https://doi.org/10.1111/evo.13075

Bieri, P., Leibundgut, M., Saurer, M., Boehringer, D., & Ban, N. (2017). The complete structure of the chloroplast 70S ribosome in complex with translation factor pY. The EMBO Journal, 36(4), 475–486. https://doi.org/10.15252/embj.201695959

Birchler, J. A., & Veitia, R. A. (2012). Gene balance hypothesis: Connecting issues of dosage sensitivity across biological disciplines. Proceedings of the National Academy of Sciences of the United States of America, 109(37), 14746–14753. https://doi.org/10.1073/pnas.1207726109

Boerema, A. P., Aibara, S., Paul, B., Tobiasson, V., Kimanius, D., Forsberg, B. O., Wallden, K., Lindahl, E., & Amunts, A. (2018). Structure of the chloroplast ribosome with chl-RRF and hibernation-promoting factor. Nature Plants, 4(4), 212–217. https://doi.org/10.1038/s41477-018-0129-6

Bogdanova, V. S., Zaytseva, O. O., Mglinets, A. V., Shatskaya, N. V., Kosterin, O. E., & Vasiliev, G. V. (2015). Nuclear-cytoplasmic conflict in pea (Pisum sativum L.) is associated with nuclear and plastidic candidate genes encoding acetyl-coA carboxylase subunits. PLoS ONE, 10(3), 1–18. https://doi.org/10.1371/journal.pone.0119835

Bonen, L., & Calixte, S. (2005). Comparative aqnalysis of bacterial-origin genes for plant mitochondrial ribosomal proteins. Mol Biol Evol, 23(3), 701–712. https://doi.org/10.1093/molbev/msj080

Castresana, J. (2000). Selection of conserved blocks from multiple alignments for their use in phylogenetic analysis. Molecular Biology and Evolution, 17(4), 540–552. https://doi.org/10.1093/oxfordjournals.molbev.a026334

Cho, Y., Mower, J. P., Qiu, Y. L., & Palmer, J. D. (2004). Mitochondrial substitution rates are extraordinarily elevated and variable in a genus of flowering plants. Proceedings of the National Academy of Sciences of the United States of America, 101(51), 17741–17746. https://doi.org/10.1073/pnas.0408302101

Christian, R. W., Hewitt, S. L., Roalson, E. H., & Dhingra, A. (2020). Genome-Scale Characterization of Predicted Plastid-Targeted Proteomes in Higher Plants. Scientific Reports, 10(1), 1–22. https://doi.org/10.1038/s41598-020-64670-5

Clark, Nathan L, Alani, E., & Aquadro, C. F. (2012). Evolutionary Rate Covariation : A bioinformatic method that reveals co-functionality and co-expression of genes. Genome Research, 22(607), 714–720. https://doi.org/10.1101/gr.132647.111.Guerois

Clark, Nathaniel L., & Aquadro, C. F. (2010). A novel method to detect proteins evolving at correlated rates: Identifying new functional relationships between coevolving proteins. Molecular Biology and Evolution, 27(5), 1152–1161. https://doi.org/10.1093/molbev/msp324

Colombo, M., Tadini, L., Peracchio, C., Ferrari, R., & Pesaresi, P. (2016). GUN1, a jack-of-all-trades in chloroplast protein homeostasis and signaling. Frontiers in Plant Science, 7(September). https://doi.org/10.3389/fpls.2016.01427

Costello, R., Emms, D. M., & Kelly, S. (2020). Gene duplication accelerates the pace of protein gain and loss from plant organelles. Molecular Biology and Evolution, 37(4), 969–981. https://doi.org/10.1093/molbev/msz275

Couvillion, M. T., Soto, I. C., Shipkovenska, G., & Churchman, L. S. (2016). Synchronized mitochondrial and cytosolic translation programs. Nature, 533(7604), 499–503. https://doi.org/10.1038/nature18015

Creff, A., Sormani, R., & Desnos, T. (2010). The two Arabidopsis RPS6 genes, encoding for cytoplasmic ribosomal proteins S6, are functionally equivalent. Plant Molecular Biology, 73(4), 533–546. https://doi.org/10.1007/s11103-010-9639-y

De Juan, D., Pazos, F., & Valencia, A. (2013). Emerging methods in protein co-evolution. Nature Reviews Genetics, 14(4), 249–261. https://doi.org/10.1038/nrg3414

De Smet, R., Adams, K. L., Vandepoele, K., Van Montagu, M. C. E., Maere, S., & Van de Peer, Y. (2013). Convergent gene loss following gene and genome duplications creates single-copy families in flowering plants. Proceedings of the National Academy of Sciences of the United States of America, 110(8), 2898–2903. https://doi.org/10.1073/pnas.1300127110/-/DCSupplemental.www.pnas.org/cgi/doi/10.1073/pnas.1300127110

Degnan, J. H., & Rosenberg, N. a. (2009). Gene tree discordance, phylogenetic inference and the multispecies coalescent. Trends in Ecology & Evolution, 24(6), 332–340. https://doi.org/10.1016/j.tree.2009.01.009

Dogra, V., Duan, J., Lee, K. P., & Kim, C. (2019). Impaired PSII proteostasis triggers a UPR-like response in the var2 mutant of Arabidopsis. Journal of Experimental Botany, 70(12), 3075–3088. https://doi.org/10.1093/jxb/erz151

Duarte, J. M., Wall, P. K., Edger, P. P., Landherr, L. L., Ma, H., Pires, J. C., Leebens-Mack, J., & dePamphilis, C. W. (2010). Identification of shared single copy nuclear genes in Arabidopsis, Populus, Vitis and Oryza and their phylogenetic utility across various taxonomic levels. BMC Evolutionary Biology, 10, 61. https://doi.org/10.1186/1471-2148-10-61

Dugas, D. V., Hernandez, D., Koenen, E. J. M., Schwarz, E., Straub, S., Hughes, C. E., Jansen, R. K., Nageswara-Rao, M., Staats, M., Trujillo, J. T., Hajrah, N. H., Alharbi, N. S., Al-Malki, A. L., Sabir, J. S. M., & Bailey, C. D. (2015). Mimosoid legume plastome evolution: IR expansion, tandem repeat expansions, and accelerated rate of evolution in clpP. Scientific Reports, 5(October), 1–13. https://doi.org/10.1038/srep16958

Emms, D. M., & Kelly, S. (2015). OrthoFinder: solving fundamental biases in whole genome comparisons dramatically improves orthogroup inference accuracy. Genome Biology, 16(1), 157. https://doi.org/10.1186/s13059-015-0721-2

Felsenstein, J. (1985). Phylogenies and the comparative method. American Naturalist, 125(1), 1–15. https://doi.org/10.1086/284325

Ferro, M., Brugière, S., Salvi, D., Seigneurin-Berny, D., Court, M., Moyet, L., Ramus, C., Miras, S., Mellal, M., Le Gall, S., Kieffer-Jaquinod, S., Bruley, C., Garin, J., Joyard, J., Masselon, C., & Rolland, N. (2010). AT-CHLORO, a comprehensive chloroplast proteome database with subplastidial localization and curated information on envelope proteins. Molecular and Cellular Proteomics, 9(6), 1063–1084. https://doi.org/10.1074/mcp.M900325-MCP200

Ferro, M., Salvi, D., Brugière, S., Miras, S., Kowalski, S., Louwagie, M., Garin, J., Joyard, J., & Rolland, N. (2003). Proteomics of the chloroplast envelope membranes from Arabidopsis thaliana. Molecular & Cellular Proteomics : MCP, 2(5), 325–345. https://doi.org/10.1074/mcp.M300030-MCP200

Findlay, G. D., Sitnik, J. L., Wang, W., Aquadro, C. F., Clark, N. L., & Wolfner, M. F. (2014). Evolutionary Rate Covariation Identifies New Members of a Protein Network Required for Drosophila melanogaster Female Post-Mating Responses. PLoS Genetics, 10(1). https://doi.org/10.1371/journal.pgen.1004108

Forsythe, E. S., Nelson, A. D. L., & Beilstein, M. A. (2020). Biased gene retention in the face of introgression obscures species relationships. Genome Biology and Evolution, 1–44.

Forsythe, E. S., Sharbrough, J., Havird, J. C., Warren, J. M., & Sloan, D. B. (2019). CyMIRA: The Cytonuclear Molecular Interactions Reference for Arabidopsis. Genome Biology and Evolution, 11(8), 2194–2202. https://doi.org/10.1093/gbe/evz144

Goh, C. S., Bogan, A. A., Joachimiak, M., Walther, D., & Cohen, F. E. (2000). Co-evolution of proteins with their interaction partners. Journal of Molecular Biology, 299(2), 283–293. https://doi.org/10.1006/jmbi.2000.3732

Gould, S. B., Waller, R. F., & McFadden, G. I. (2008). Plastid Evolution. Annual Review of Plant Biology, 59(1), 491–517. https://doi.org/10.1146/annurev.arplant.59.032607.092915

Greiner, S., Rauwolf, U., Meurer, J., & Herrmann, R. G. (2011). The role of plastids in plant speciation. Molecular Ecology, 20(4), 671–691. https://doi.org/10.1111/j.1365-294X.2010.04984.x

Guisinger, M. M., Kuehl, J. V., Boore, J. L., & Jansen, R. K. (2008). Genome-wide analyses of Geraniaceae plastid DNA reveal unprecedented patterns of increased nucleotide substitutions. Proceedings of the National Academy of Sciences of the United States of America, 105(47), 18424–18429. https://doi.org/10.1073/pnas.0806759105

Guisinger, M. M., Kuehl, J. V., Boore, J. L., & Jansen, R. K. (2011). Extreme reconfiguration of plastid genomes in the angiosperm family Geraniaceae: Rearrangements, repeats, and codon usage. Molecular Biology and Evolution, 28(1), 583–600. https://doi.org/10.1093/molbev/msq229

Hilu, K. W., Borsch, T., Müller, K., Soltis, D. E., Soltis, P. S., Savolainen, V., Chase, M. W., Powell, M. P., Alice, L. A., Evans, R., Sauquet, H., Neinhuis, C., Slotta, T. A. B., Rohwer, J. G., Campbell, C. S., & Chatrou, L. W. (2003). Angiosperm phylogeny based on matK sequence information. American Journal of Botany, 90(12), 1758–1776. https://doi.org/10.3732/ajb.90.12.1758

Houtkooper, R. H., Mouchiroud, L., Ryu, D., Moullan, N., Katsyuba, E., Knott, G., Williams, R. W., & Auwerx, J. (2013). Mitonuclear protein imbalance as a conserved longevity mechanism. Nature, 497(7450), 451–457. https://doi.org/10.1038/nature12188

Jansen, R. K., Cai, Z., Raubeson, L. A., Daniell, H., Depamphilis, C. W., Leebens-Mack, J., Müller, K. F., Guisinger-Bellian, M., Haberle, R. C., Hansen, A. K., Chumley, T. W., Lee, S. B., Peery, R., McNeal, J. R., Kuehl, J. V., & Boore, J. L. (2007). Analysis of 81 genes from 64 plastid genomes resolves relationships in angiosperms and identifies genome-scale evolutionary patterns. Proceedings of the National Academy of Sciences of the United States of America, 104(49), 19369–19374. https://doi.org/10.1073/pnas.0709121104

Katoh, K., & Standley, D. M. (2013). MAFFT multiple sequence alignment software version 7: Improvements in performance and usability. Molecular Biology and Evolution, 30(4), 772–780. https://doi.org/10.1093/molbev/mst010

Kikuchi, S., Asakura, Y., Imai, M., Nakahira, Y., Kotani, Y., Hashiguchi, Y., Nakai, Y., Takafuji, K., Bédard, J., Hirabayashi-Ishioka, Y., Mori, H., Shiina, T., & Nakai, M. (2018). A Ycf2-FtsHi heteromeric AAA-ATPase complex is required for chloroplast protein import. Plant Cell, 30(11), 2677–2703. https://doi.org/10.1105/tpc.18.00357

Kim, J., Olinares, P. D., Oh, S. H., Ghisaura, S., Poliakov, A., Ponnala, L., & van Wijk, K. J. (2013). Modified Clp protease complex in the ClpP3 null mutant and consequences for chloroplast development and function in Arabidopsis. Plant Physiology, 162(1), 157–179. https://doi.org/10.1104/pp.113.215699

Knox, E. B. (2014). The dynamic history of plastid genomes in the Campanulaceae sensu lato is unique among angiosperms. Proceedings of the National Academy of Sciences of the United States of America, 111(30), 11097–11102. https://doi.org/10.1073/pnas.1403363111

Leebens-Mack, Barker, Carpenter, et al. (2019). One thousand plant transcriptomes and the phylogenomics of green plants. 16(November 2017). https://doi.org/10.1038/s41586-019-1693-2

Liu, X., Yu, F., & Rodermel, S. (2010). An arabidopsis pentatricopeptide repeat protein, SUPPRESSOR OF VARIEGATION7, is required for FtsH–mediated chloroplast biogenesis. Plant Physiology, 154(4), 1588–1601. https://doi.org/10.1104/pp.110.164111

Magee, A. M., Aspinall, S., Rice, D. W., Cusack, B. P., Sémon, M., Perry, A. S., Stefanović, S., Milbourne, D., Barth, S., Palmer, J. D., Gray, J. C., Kavanagh, T. A., & Wolfe, K. H. (2010). Localized hypermutation and associated gene losses in legume chloroplast genomes. Genome Research, 20(12), 1700–1710. https://doi.org/10.1101/gr.111955.110

Millar, A. H., Whelan, J., & Small, I. (2006). Recent surprises in protein targeting to mitochondria and plastids. Current Opinion in Plant Biology, 9(6), 610–615. https://doi.org/10.1016/j.pbi.2006.09.002

Molina, J., Hazzouri, K. M., Nickrent, D., Geisler, M., Meyer, R. S., Pentony, M. M., Flowers, J. M., Pelser, P., Barcelona, J., Inovejas, S. A., Uy, I., Yuan, W., Wilkins, O., Michel, C. I., Locklear, S., Concepcion, G. P., & Purugganan, M. D. (2014). Possible loss of the chloroplast genome in the parasitic flowering plant Rafflesia lagascae (Rafflesiaceae). Molecular Biology and Evolution, 31(4), 793–803. https://doi.org/10.1093/molbev/msu051

Mower, J. P., Touzet, P., Gummow, J. S., Delph, L. F., & Palmer, J. D. (2007). Extensive variation in synonymous substitution rates in mitochondrial genes of seed plants. BMC Evolutionary Biology, 7, 1–14. https://doi.org/10.1186/1471-2148-7-135

Nevill, P. G., Howell, K. A., Cross, A. T., Williams, A. V., Zhong, X., Tonti-Filippini, J., Boykin, L. M., Dixon, K. W., & Small, I. (2019). Plastome-wide rearrangements and gene losses in carnivorous droseraceae. Genome Biology and Evolution, 11(2), 472–485. https://doi.org/10.1093/gbe/evz005

Nishimura, K., & van Wijk, K. J. (2015). Organization, function and substrates of the essential Clp protease system in plastids. BBA -Bioenergetics, 1847(9), 915–930. https://doi.org/10.1016/j.bbabio.2014.11.012

Panchy, N., Lehti-Shiu, M., & Shiu, S. H. (2016). Evolution of gene duplication in plants. Plant Physiology, 171(4), 2294–2316. https://doi.org/10.1104/pp.16.00523

Paradis, E., Claude, J., & Strimmer, K. (2004). APE: Analyses of phylogenetics and evolution in R language. Bioinformatics, 20(2), 289–290. https://doi.org/10.1093/bioinformatics/btg412

Park, Seongjun, Ruhlman, T. A., Weng, M. L., Hajrah, N. H., Sabir, J. S. M., & Jansen, R. K. (2017). Contrasting patterns of nucleotide substitution rates provide insight into dynamic evolution of plastid and mitochondrial genomes of Geranium. Genome Biology and Evolution, 9(6), 1766–1780. https://doi.org/10.1093/gbe/evx124

Park, Sungsoon, & Rodermel, S. R. (2004). Mutations in ClpC2/Hsp100 suppress the requirement for FtsH in thylakoid membrane biogenesis. Proceedings of the National Academy of Sciences of the United States of America, 101(34), 12765–12770. https://doi.org/10.1073/pnas.0402764101

Parkinson, C. L., Mower, J. P., Qiu, Y. L., Shirk, A. J., Song, K., Young, N. D., DePamphilis, C. W., & Palmer, J. D. (2005). Multiple major increases and decreases in mitochondrial substitution rates in the plant family Geraniaceae. BMC Evolutionary Biology, 5, 1–12. https://doi.org/10.1186/1471-2148-5-73

Qiu, Y. L., Dombrovska, O., Lee, J., Li, L., Whitlock, B. A., Bernasconi-Quadroni, F., Rest, J. S., Davis, C. C., Borsch, T., Hilu, K. W., Renner, S. S., Soltis, D. E., Soltis, P. S., Zanis, M. J., Cannone, J. J., Gutell, R. R., Powell, M., Savolainen, V., Chatrou, L. W., & Chase, M. W. (2005). Phylogenetic analyses of basal angiosperms based on nine plastid, mitochondrial, and nuclear genes. International Journal of Plant Sciences, 166(5), 815–842. https://doi.org/10.1086/431800

Qiu, Y. L., Li, L., Hendry, T. A., Li, R., Taylor, D. W., Issa, M. J., Ronen, A. J., Vekaria, M. L., & White, A. M. (2006). Reconstructing the basal angiosperm phylogeny: Evaluating information content of mitochondrial genes. Taxon, 55(4), 837–856. https://doi.org/10.2307/25065680

Ramani, A. K., & Marcotte, E. M. (2003). Exploiting the co-evolution of interacting proteins to discover interaction specificity. Journal of Molecular Biology, 327(1), 273–284. https://doi.org/10.1016/S0022-2836(03)00114-1

Raza, Q., Choi, J. Y., Li, Y., O’Dowd, R. M., Watkins, S. C., Chikina, M., Hong, Y., Clark, N. L., & Kwiatkowski, A. V. (2019). Evolutionary rate covariation analysis of E-cadherin identifies Raskol as a regulator of cell adhesion and actin dynamics in Drosophila. PLoS Genetics, 15(2), 1–24. https://doi.org/10.1371/journal.pgen.1007720

Rockenbach, K., Havird, J. C., Grey Monroe, J., Triant, D. A., Taylor, D. R., & Sloan, D. B. (2016). Positive selection in rapidly evolving plastid-nuclear enzyme complexes. Genetics, 204(4), 1507–1522. https://doi.org/10.1534/genetics.116.188268

Sanderson, M. J., & McMahon, M. M. (2007). Inferring angiosperm phylogeny from EST data with widespread gene duplication. BMC Evolutionary Biology, 7 Suppl 1, S3. https://doi.org/10.1186/1471-2148-7-S1-S3

Sangiovanni, M., Vigilante, A., & Chiusano, M. (2013). Exploiting a Reference Genome in Terms of Duplications: The Network of Paralogs and Single Copy Genes in Arabidopsis thaliana. Biology, 2(4), 1465–1487. https://doi.org/10.3390/biology2041465

Sato, T., Yamanishi, Y., Kanehisa, M., & Toh, H. (2005). The inference of protein-protein interactions by co-evolutionary analysis is improved by excluding the information about the phylogenetic relationships. Bioinformatics, 21(17), 3482–3489. https://doi.org/10.1093/bioinformatics/bti564

Schmitz-Linneweber, C., Kushnir, S., Babiychuk, E., Poltnigg, P., Herrmann, R. G., & Maier, R. M. (2005). Pigment deficiency in nightshade/tobacco cybrids is caused by the failure to edit the plastid ATPase α-subunit mRNA. Plant Cell, 17(6), 1815–1828. https://doi.org/10.1105/tpc.105.032474

Sharbrough, J., Conover, J. L., Tate, J. A., Wendel, J. F., & Sloan, D. B. (2017). Cytonuclear responses to genome doubling. 104(9), 1–4. https://doi.org/10.3732/ajb.1700293

Shrestha, B., Weng, M. L., Theriot, E. C., Gilbert, L. E., Ruhlman, T. A., Krosnick, S. E., & Jansen, R. K. (2019). Highly accelerated rates of genomic rearrangements and nucleotide substitutions in plastid genomes of Passiflora subgenus Decaloba. Molecular Phylogenetics and Evolution, 138(May), 53–64. https://doi.org/10.1016/j.ympev.2019.05.030

Sloan, D. B., Oxelman, B., Rautenberg, A., & Taylor, D. R. (2009). Phylogenetic analysis of mitochondrial substitution rate variation in the angiosperm tribe Sileneae. BMC Evolutionary Biology, 9(1), 1–16. https://doi.org/10.1186/1471-2148-9-260

Sloan, D. B., Triant, D. A., Forrester, N. J., Bergner, L. M., Wu, M., & Taylor, D. R. (2014). A recurring syndrome of accelerated plastid genome evolution in the angiosperm tribe Sileneae (Caryophyllaceae). Molecular Phylogenetics and Evolution, 72(1), 82–89. https://doi.org/10.1016/j.ympev.2013.12.004

Sloan, D. B., Triant, D. A., Wu, M., & Taylor, D. R. (2014). Cytonuclear interactions and relaxed selection accelerate sequence evolution in organelle ribosomes. Molecular Biology and Evolution, 31(3), 673–682. https://doi.org/10.1093/molbev/mst259

Smith, M. L., & Hahn, M. W. (2020). New approaches for inferring phylogenies in the presence of paralogs. BioRxiv, 1, 6–8. https://doi.org/10.16309/j.cnki.issn.1007-1776.2003.03.004

Sobanski, J., Giavalisco, P., Fischer, A., Kreiner, J. M., Walther, D., Aurel, M., Pellizzer, T., Golczyk, H., Obata, T., Bock, R., Sears, B. B., & Greiner, S. (2019). Chloroplast competition is controlled by lipid biosynthesis in evening primroses. 116(12). https://doi.org/10.1073/pnas.1811661116

Soltis, P. S., Soltis, D. E., & Chase, M. W. (1999). Angiosperm phylogeny inferred from multiple genes as a tool for comparative biology. Nature, 402(6760), 402–404. https://doi.org/10.1038/46528

Stamatakis, A. (2014). RAxML version 8: a tool for phylogenetic analysis and post-analysis of large phylogenies. Bioinformatics (Oxford, England), 30(9), 1312–1313. https://doi.org/10.1093/bioinformatics/btu033

Stolzer, M., Lai, H., Xu, M., Sathaye, D., Vernot, B., & Durand, D. (2012). Inferring duplications, losses, transfers and incomplete lineage sorting with nonbinary species trees. Bioinformatics, 28(18), i409–i415. https://doi.org/10.1093/bioinformatics/bts386

Tiller, N., Weingartner, M., Thiele, W., Maximova, E., & Scho, M. A. (2012). The plastid-specific ribosomal proteins of Arabidopsis thaliana can be divided into non-essential proteins and genuine ribosomal proteins. The Plan Journal, 69, 302–316. https://doi.org/10.1111/j.1365-313X.2011.04791.x

Timmis, J. N., Ayliff, M. A., Huang, C. Y., & Martin, W. (2004). Endosymbiotic gene transfer: Organelle genomes forge eukaryotic chromosomes. Nature Reviews Genetics, 5(2), 123–135. https://doi.org/10.1038/nrg1271

van Wijk, K. J., & Baginsky, S. (2011). Plastid proteomics in higher plants: Current state and future goals. Plant Physiology, 155(4), 1578–1588. https://doi.org/10.1104/pp.111.172932

Vernot, B., Stolzer, M., Goldman, A., & Durand, D. (2008). Reconciliation with non-binary species trees. Journal of Computational Biology : A Journal of Computational Molecular Cell Biology, 15(8), 981–1006. https://doi.org/10.1089/cmb.2008.0092

Wagner, R., Aigner, H., & Funk, C. (2012). FtsH proteases located in the plant chloroplast. Physiologia Plantarum, 145(1), 203–214. https://doi.org/10.1111/j.1399-3054.2011.01548.x

Waltz, F., Nguyen, T. T., Arrivé, M., Bochler, A., Chicher, J., Hammann, P., Kuhn, L., Quadrado, M., Mireau, H., Yashem, Y., & Giegé, P. (2019). Small is big in Arabidopsis mitochondrial ribosome. Nature Plants, 5(January), 106–117. https://doi.org/10.1038/s41477-018-0339-y

Wang, R., Zhao, J., Jia, M., Xu, N., Liang, S., Shao, J., Qi, Y., Liu, X., An, L., & Yu, F. (2018). Balance between cytosolic and chloroplast translation affects leaf variegation. Plant Physiology, 176(1), 804–818. https://doi.org/10.1104/pp.17.00673

Wendel, J. F., Lisch, D., Hu, G., & Mason, A. S. (2018). The long and short of doubling down: polyploidy, epigenetics, and the temporal dynamics of genome fractionation. Current Opinion in Genetics and Development, 49, 1–7. https://doi.org/10.1016/j.gde.2018.01.004

Weng, M., Ruhlman, T. A., & Jansen, R. K. (2016). Plastid – Nuclear Interaction and Accelerated Coevolution in Plastid Ribosomal Genes in Geraniaceae. 8(6), 1824–1838. https://doi.org/10.1093/gbe/evw115

Wicke, S., Müller, K. F., DePamphilis, C. W., Quandt, D., Bellot, S., & Schneeweiss, G. M. (2016). Mechanistic model of evolutionary rate variation en route to a nonphotosynthetic lifestyle in plants. Proceedings of the National Academy of Sciences of the United States of America, 113(32), 9045–9050. https://doi.org/10.1073/pnas.1607576113

Williams, A. M., Friso, G., Wijk, K. J. Van, & Sloan, D. B. (2019). Extreme variation in rates of evolution in the plastid Clp protease complex. The Plan Journal, 1–17. https://doi.org/10.1111/tpj.14208

Wolfe, N. W., & Clark, N. L. (2015). ERC analysis: Web-based inference of gene function via evolutionary rate covariation. Bioinformatics, 31(23), 3835–3837. https://doi.org/10.1093/bioinformatics/btv454

Yan, Z., Ye, G., & Werren, J. H. (2019). Evolutionary Rate Correlation between Mitochondrial-Encoded and Mitochondria-Associated Nuclear-Encoded Proteins in Insects. Molecular Biology and Evolution, 36(5), 1022–1036. https://doi.org/10.1093/molbev/msz036

Yu, F., Liu, X., Alsheikh, M., Park, S., & Rodermel, S. (2008). Mutations in SUPPRESSOR OF VARIEGATION1, a factor required for normal chloroplast translation, suppress var2-mediated leaf variegation in Arabidopsis. The Plant Cell, 20(July), 1786–1804. https://doi.org/10.1105/tpc.107.054965

Zanis, M. J., Soltis, D. E., Soltis, P. S., Mathews, S., & Donoghue, M. J. (2002). The root of the angiosperms revisited. Proceedings of the National Academy of Sciences of the United States of America, 99(10), 6848–6853. https://doi.org/10.1073/pnas.092136399

Zhang, J., Ruhlman, T. A., Sabir, J., Blazier, J. C., & Jansen, R. K. (2015). Coordinated rates of evolution between interacting plastid and nuclear genes in geraniaceae. The Plant Cell, 27(March), 563–573. https://doi.org/10.1105/tpc.114.134353

Zhang, J., Ruhlman, T. A., Sabir, J. S. M., Blazier, J. C., Weng, M., Park, S., & Jansen, R. K. (2016). Coevolution between Nuclear-Encoded DNA Replication, Recombination, and Repair Genes and Plastid Genome Complexity. 8(3), 622–634. https://doi.org/10.1093/gbe/evw033

Zheng, M., Liu, X., Liang, S., Fu, S., Qi, Y., Zhao, J., Shao, J., An, L., & Yu, F. (2016). Chloroplast translation initiation factors regulate leaf variegation and development. Plant Physiology, 172(2), 1117–1130. https://doi.org/10.1104/pp.15.02040

Zupoka, A., Kozula, D., Schöttlera, M. A., Niehörstera, J., Garbscha, F., Liereb, K., Malinovaa, I., Bock, R., & Greiner, S. (2020). A photosynthesis operon in the chloroplast genome drives speciation in evening primroses. BioRvix.

